# Vigilance, arousal, and acetylcholine: Optimal control of attention in a simple detection task

**DOI:** 10.1101/2022.02.20.481204

**Authors:** Sahiti Chebolu, Peter Dayan, Kevin Lloyd

## Abstract

Paying attention to particular aspects of the world or being more vigilant in general can be interpreted as forms of ‘internal’ action. Such arousal-related choices come with the benefit of increasing the quality and situational appropriateness of information acquisition and processing, but incur potentially expensive energetic and opportunity costs. One implementational route for these choices is widespread ascending neuromodulation, including by acetylcholine (ACh). The key computational question that elective attention poses for sensory processing is when it is worthwhile paying these costs, and this includes consideration of whether sufficient information has yet been collected to justify the higher signal-to-noise ratio afforded by greater attention and, particularly if a change in attentional state is more expensive than its maintenance, when states of heightened attention ought to persist. We offer a partially observable Markov decision-process treatment of optional attention in a detection task, and use it to provide a qualitative model of the results of studies using modern techniques to measure and manipulate ACh in rodents performing a similar task.

## 1 Introduction

*Vigilance* is commonly taken to mean the state of being alertly watchful, and is principally equated with sustained attention in the scientific literature (Davies & Parasuraman, 1982; Oken, Salinsky, & Elsas, 2006). People notoriously find it difficult and taxing to maintain high levels of vigilance, or sustained attention, over long periods when required by the exigencies of a task, typically exhibiting fluctuations in their performance as well as an overall decline in performance over time (reviews include Fortenbaugh, DeGutis, & Esterman, 2017; Gilden & Wilson, 1995; Warm, Parasuraman, & Matthews, 2008). Vigilance was particularly actively investigated in the middle of the last century, for instance in the case of radar operators having to search for targets over long periods (Broadbent, 1971; N. Mackworth, 1948), and was commonly related to *arousal*, which was generally conceptualized in more physiological terms as the level of non-specific activation of cerebral cortex (Hebb, 1955; Lindsley, 1951; J. Mackworth, 1968; though see Duffy, 1957). Vigilance has also been explored in non-human animals, such as in the sustained attention task developed for rodents by McGaughy and Sarter (1995).

However, rather like selective forms of attention (Dayan & Solomon, 2010; Dayan & Zemel, 1999), there remain gaps in the examination of vigilance using modern conceptions of cost-sensitive information-processing in the brain, as, for instance, in the framework of the expected value of control (EVC; Shenhav, Botvinick, & Cohen, 2013; Shenhav et al., 2017; though see the recent review by Esterman & Rothlein, 2019), in which the degree of cognitive control applied in a situation is normatively determined by a cost-benefit analysis. Thus, in the EVC framework, putative benefits of vigilance might include an increased signal-to-noise ratio (SNR) for sensation and cognition, but at the expense of actual costs of excess neural activity and/or opportunity costs from the diversion of processing to the task at hand rather than other potentially valuable internally- or externally-focused computations (Boureau, Sokol-Hessner, & Daw, 2015; Esterman & Rothlein, 2019; Kurzban, Duckworth, Kable, & Myers, 2013; Warm et al., 2008).

Concomitantly, EVC conceptions could be enriched by insights from the human and animal paradigms concerned with the neural foundations of vigilance, which, for instance, point to the central involvement of neuromodulatory systems (Aston-Jones & Cohen, 2005; Oken et al., 2006; Parasuraman, Warm, & See, 1998; Robbins, 1997, 2002). Neuromodulators can be seen as one solution adopted by the brain for centralised regulation of distributed processing (Dayan, 2012b). This has at least two facets: increasing excitability to boost the SNR of neural representations; and, since vigilance is necessary when aspects of the external environment are known to be unknown, manipulating the balance between the influence of prior expectations versus input likelihoods. The latter is the putative role of ACh as the medium of expected uncertainty in the theoretical suggestion of Yu and Dayan (2002, 2005).

Here, we use an abstract model of the sustained attention task (SAT) of McGaughy and Sarter (1995) to elaborate the statistical and computational bases of vigilance in EVC terms, and explore qualitative aspects of the role of neuromodulation — specifically, ACh — in controlled information-processing. Thus, we note at the outset that our study focuses on one particular experimental paradigm, and on findings about ACh gleaned from that paradigm; this means that we certainly do not claim to address all of the broad range of findings on vigilance more generally, which also frequently involve rather different tasks (see, e.g., Fortenbaugh et al., 2017).

In the SAT, animals face repeated trials in each of which they have to watch out for a potentially extremely brief signal (a flash of light at a particular location) that comes at an uncertain time during an extended epoch. The end of this epoch is indicated by the insertion of two levers into the operant chamber, and the animal indicates whether it believes a signal has been present or absent during the trial by its selection of lever. Correct responses (i.e. hits and correct rejections) are rewarded, while incorrect responses (i.e., misses and false alarms) are unrewarded. Withdrawal of the levers, either once the animal has made a response or after the maximum response time of 4 s, marks the beginning of the next trial. Uncertainty about whether and when the brief signal will arrive means the animals have to be vigilant across the whole epoch. Many studies (Gritton et al., 2016; Hasselmo & Sarter, 2011; McGaughy, Dalley, Morrison, Everitt, & Robbins, 2002; McGaughy, Kaiser, & Sarter, 1996; McGaughy & Sarter, 1998; Sarter & Lustig, 2019) have shown that good quality performance depends causally on cholinergic neuromodulation associated both with basal forebrain sources of ACh and controlled release in the medial prefrontal cortex; complex patterns of ACh release over sequences of trials have also been observed (Howe et al., 2013). This long line of investigation by Sarter and colleagues into the role of ACh in the SAT make it a natural target for computational modelling.

From a computational perspective, we treat the decision problem faced by an animal in the SAT as a form of partially observed Markov decision process (POMDP; Kaelbling, Littman, & Cassandra, 1998). We suggest that animals can be more or less vigilant (i.e., have *strong* or *weak* attention), with the difference reflected in the SNR of the input, but also with costs for switching into, and maintaining, *strong* attention (cf. Atkinson, 1963). Thus, the agent’s objective is not exactly to report correctly whether a trial contains a signal or not at the trial’s conclusion, but rather to maximize the difference between external rewards (provided by the experimenter based on successful task execution) and internal costs (from the engagement of attention over the course of the trial) by appropriate choice of both attentional states and final report. The existence of different attentional states (*strong* vs. *weak*) and the ability to control which attentional state is occupied makes for a form of extended signal detection theory problem with explicitly controlled variable signal quality (Cisek, Puskas, & El-Murr, 2009; Drugowitsch, Moreno-Bote, & Pouget, 2014; Hébert & Woodford, 2017; Jang, Sharma, & Drugowitsch, 2021). Our computational agent has to use the information it has acquired during a trial so far to decide whether it is worth paying the cost of switching from *weak* to *strong* attention. If so, then consistent with the sustained attention styling of the task, it has to decide whether it is worth maintaining the state of elevated attention across successive trials.

We interpret the attentional switching as an internal action (Dayan, 2012a) mediated by ACh (albeit itself under the influence of extensive cortical control), and use our model to interpret the neuromodulatory findings of Gritton et al. (2016); Howe et al. (2013) in terms of a phasic theory of ACh, to complement the tonic characterization offered in Yu and Dayan (2005). One important difference from the tonic theory is that there is no exact equivalence between *weak*/*strong* and top-down/bottom-up information influences in the cortex. Rather, we treat *strong* attention as a top-down controlled state in which bottom-up information is more reliable (and typically known to be more reliable — the consequences of possible mismatches between what is actual and what is known or believed is considered in Section 3.2, below), and so exerts a greater influence over hierarchical processing.

We first describe the abstraction of the SAT, and then consider the deployment of attention over the course of single trials, showing conditions under which it is adaptive to switch from *weak* to *strong* attention when a sufficient hint of a signal is detected. We show that even such a simple model of attention can lead to rich patterns of behaviour, both within and across trials, and in a manner that depends in subtle ways on expectations about whether and when a signal may appear, informational quality, and assumed attentional costs. We also report subtle issues concerning the source of misses and false alarms when ACh is manipulated (Gritton et al., 2016; McGaughy et al., 1996). We next consider sustained attention across multiple trials, particularly in the light of findings about specific circumstances under which ACh is engaged in the task (Howe et al., 2013). Finally, we discuss our results and interpretations in the light of EVC, expected uncertainty, and older ideas about vigilance and arousal.

## 2 Methods

### 2.1 Model

Briefly, we model an abstract version of the SAT (Fig. 1A) in which a potentially very brief signal may or may not occur (and if so, at an uncertain time) during an extended period. At the end of each trial, the decision-maker reports whether or not they think a signal occurred during the trial. Imperfect information about signal presence or absence is carried by noisy observations. Crucially, how informative these observations are depends on the attentional state of the agent, which we assume to be under its direct control. On each time step, the agent makes a binary choice about whether to attend *weakly* or *strongly*– strong attention yields better quality information, but at a cost. We consider the problem of how to optimize this sequence of decisions regarding attentional state, balancing the putative costs of enhanced attention against the benefits of better evidence. With the exception of Section 3.2, in which we consider how optogenetic manipulations may disturb correct inference, we assume that the actual and believed qualities of the signal are identical. Formally, the decision problem is modelled as a partially observed Markov decision process (POMDP; cf. Fig. 1B), with elements detailed in the following.

**Figure 1:**
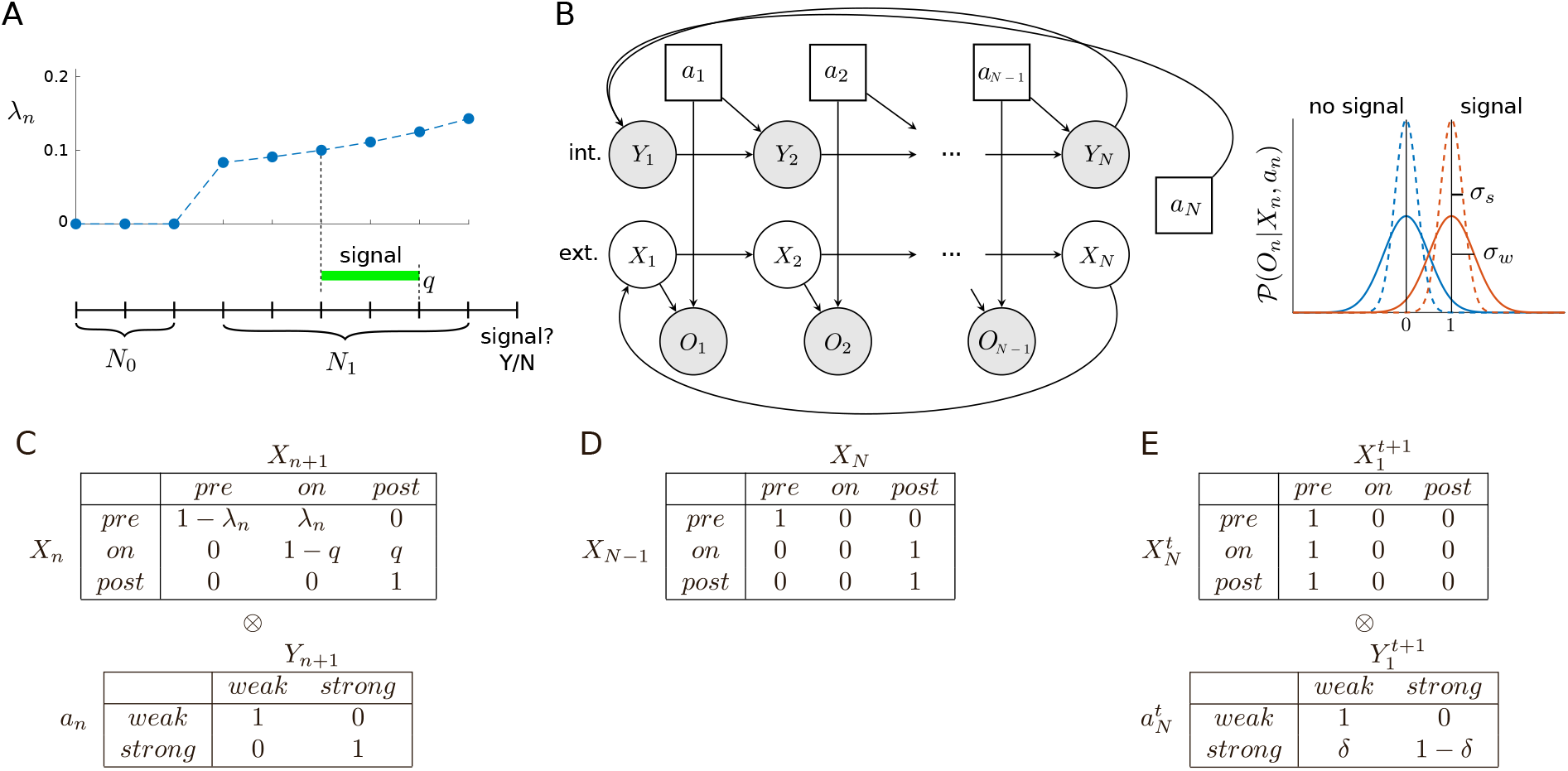
Model. (A) Each trial, in which a signal may or may not occur, comprises a sequence of *N* = *N*_0_ + *N*_1_ + 1 time steps. Whether a signal occurs, and if so, its time of onset and offset, is determined by the conditional probability of arrival, *λ*_*n*_, at each step *n*, and the (constant) probability of turning off per time step, *q*. On the final time step, the decision-maker reports whether a signal did or did not occur during the trial. (B) On each time step, the state comprises the pair of variables (*X*_*n*_, *Y*_*n*_), where *X*_*n*_ ∈ {*pre, on, post*} is the (unknown) stage within the trial, and *Y*_*n*_ ∈ {*weak, strong*} is the (known) attentional state; the attentional action *a*_*n*_ ∈ {*weak, strong*} determines the quality of the observation *O*_*n*_ by determining the amount of Gaussian noise (right), and the attentional state at the next time step (if *n < N*) or beginning of the next trial (if *n* = *N*). Unshaded nodes indicate unobserved/latent random variables (actions are assumed known). For simplicity, we have omitted the additional, ‘external’ action taken at *N* (i.e., report ‘signal’ or ‘no signal’). (C) Conditional state transition probabilities. These can be decomposed into independent ‘external’ and ‘internal’ components, *𝒫*(*S*_*n*+1_|*S*_*n*_, *a*_*n*_) = *𝒫*(*X*_*n*+1_|*X*_*n*_) ⊗ *𝒫*(*Y*_*n*+1_|*a*_*n*_). (D) Transition into the final, decision state of each trial. (E) Each trial always begins in the pre-signal state, i.e., 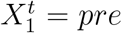 for any trial *t*; there is a probability *δ* that the next trial will be started in the *weak* attentional state even if the agent chose *strong* at the end of the previous trial.

#### 2.1.1 States

On each trial, the agent occupies a series of *N* states *S*_1_, *S*_2_, …, *S*_*N*_, where each state comprises two variables, *S*_*n*_ = (*X*_*n*_, *Y*_*n*_). *X*_*n*_ ∈ {*pre, on, post*} is an ‘external’ state component determined by the experimenter, corresponding to the three possible stages of a trial (respectively: pre-signal; signal on; and post-signal), and is not directly known by the agent; *Y*_*n*_ ∈ {*weak, strong*} is an ‘internal’ variable that reflects the agent’s current attentional state and is always assumed to be known by the agent (Fig. 1B). Formally, *Y*_*n*_ is a necessary part of the state description to allow the agent to pay different costs for switching attention from *weak* to *strong*, and for maintaining *strong* attention.

#### 2.1.2 Actions

On all time steps 1, 2, …, *N*, the agent makes a binary choice *a*_*n*_ ∈ {*weak, strong*} between whether to attend only *weakly* to sensory information, or instead to attend *strongly*. On time steps 1, 2, …, *N* − 1, choosing to attend strongly has the benefit of providing better information about the underlying state, but carries greater cost, as detailed below. At time step *N*, this same decision has no immediate informational value (since there is no observation on this step), but it determines whether the agent begins the next trial in the *weak* or *strong* attentional state — a choice of *strong* here can therefore be unambiguously interpreted as being motivated by future, rather than immediate, benefits.

In addition, on the final step *N* of each trial, the agent must indicate whether or not a signal was presented during the trial. Thus, on the final step, the agent in fact selects one of four possible actions, since it must choose both an ‘internal’, attentional action, and an ‘external’ action, i.e., *a*_*N*_ ∈ {*weak, strong*} × {*no signal, signal*}. In the following, for simplicity, *a*_*n*_ will always refer to the attentional action, unless otherwise indicated.

#### 2.1.3 State transitions

A trial contains a signal (referred to as a ‘signal trial’) with assumed (correctly or not) probability *p*_1_, and contains no signal (‘non-signal trial’) with probability *p*_0_ = 1 − *p*_1_. On all trials, we assume *N*_0_ ≥ 1 time steps where it is known that a signal cannot be present (i.e., the agent occupies *X* = *pre* with certainty), followed by *N*_1_ time steps, where a signal may arrive. We write *N* = *N*_0_ + *N*_1_ + 1, allowing for the extra time step for the {*no signal, signal*} report. A signal, should one occur, is assumed to have equal probability of arriving at any one of the *N*_1_ time steps in a trial, so that a signal’s time of arrival *τ* is

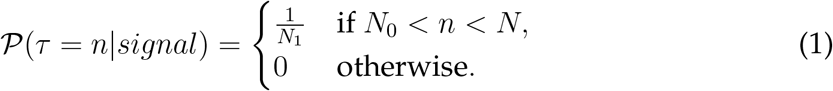

Taking into account the probability *p*_1_ that a trial contains a signal, the probability that a signal arrives on step *n* given it hasn’t arrived sooner (i.e., the hazard function) is

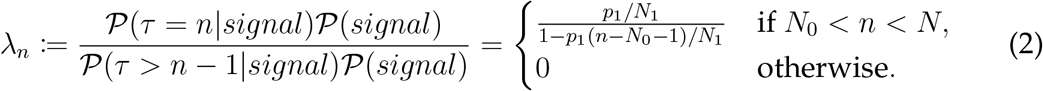

Conversely, if a signal has already arrived, we assume that it turns off again with constant probability *q* per time step (Fig. 1A). It is always turned off before the final decision step *n* = *N*.

Note that the assumption that a signal has equal probability of arriving at any one of the *N*_1_ time steps means that *λ*_*n*_ increases over the course of a trial (Fig. 2A). Note also the effect of *q* on the probability of being in the signal state at any time, *𝒫*(*X*_*n*_ = *on*).

**Figure 2:**
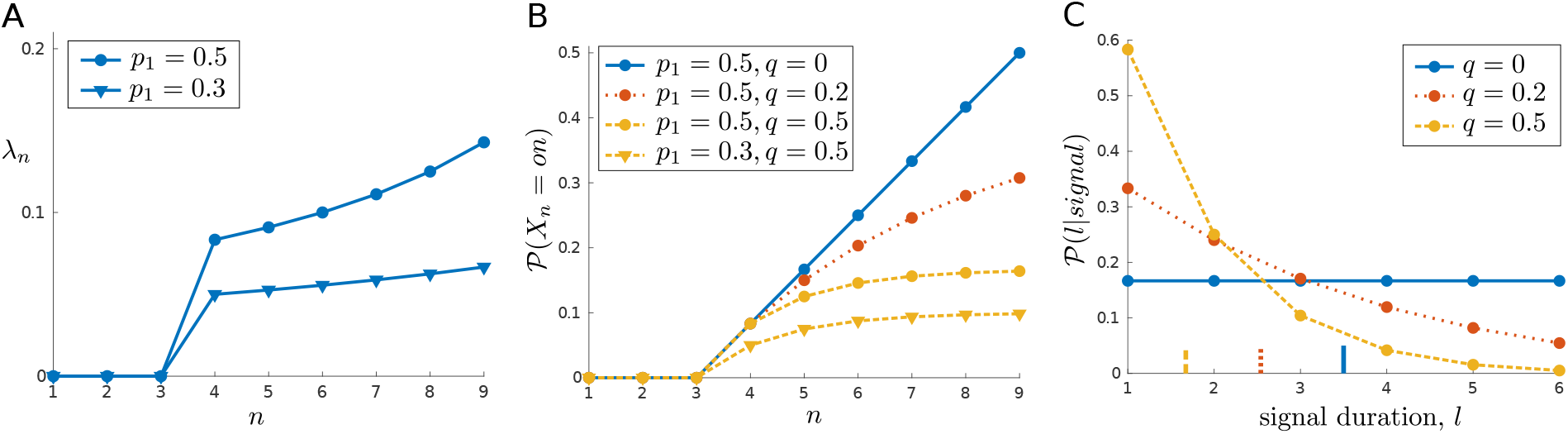
Effects of *p*_1_ and *q* on signal properties. (A) Given a uniform distribution over possible signal onset times, the conditional probability of a signal arriving given one has not yet occurred, *λ*_*n*_, increases over a trial. (B) The unconditional probability of occupying the signal *on* state at any time depends on both the prior probability of a signal, *p*_1_, and the probability of the signal turning off again once on, *q*. (C) The distribution over signal durations deviates from a geometric distribution due to the fixed trial length (e.g., mean durations, indicated by vertical lines on *x*-axis, are less than 1*/q*). In this example, *N*_0_ = 3 and *N*_1_ = 6 (so *N* = 10).

To be in the *on* state at *n*, the signal must have arrived earlier or on time step *n* and not have turned off once on. If we now let *τ*_1_ denote the time of signal onset, and *τ*_2_ denote the time of signal offset, then this probability is

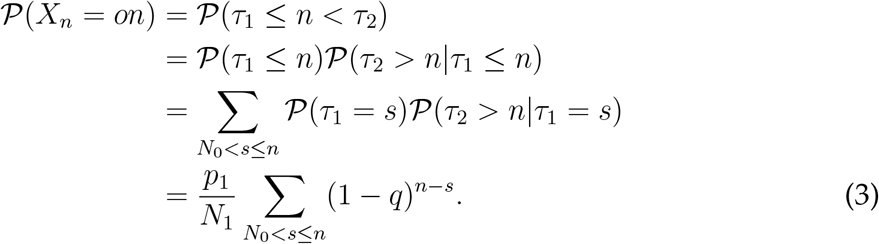

If we have *q* = 0, so that a signal would always persist until the end of the trial once on, then this probability increases to *p*_1_ just before the decision state (i.e., *𝒫*(*X*_*N*−1_ = *on*) = *p*_1_); if *q* = 1, so that a signal only ever lasts a single time step, then we have *𝒫*(*X*_*n*_ = *on*) = *p*_1_*/N*_1_ for each of the *N*_1_ time steps; and other values of *q* lead to intermediate cases (Fig. 2B). As we will see, this can affect the allocation of attention — since *strong* attention increases the detectability of a signal at a cost, it can be appropriate to wait until late in a trial when a signal is more likely to be present to be detected.

For the sake of completeness, we also note that the assumption that each trial has a fixed length of *N* steps means that the distribution over signal durations deviates, due to truncation, from a geometric distribution with parameter *q* and mean 1*/q* (Fig. 2C). The probability of a signal of duration *L* (conditioned on a signal occurring), which is nonzero only for 1, …, *N*_1_, can be expressed as

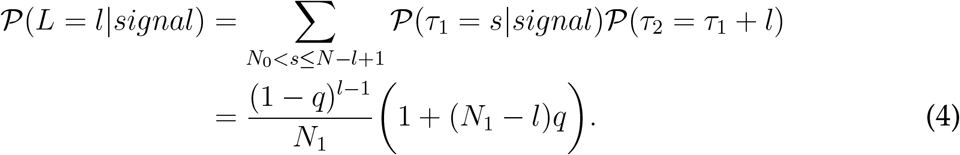

For steps 1, 2, …, *N* − 1, we assume that the attentional state on step *n*+1 is completely determined by action *a*_*n*_, so that if the choice is *weak* on step *n*, then at the start of the next time step we have *Y*_*n*+1_ = *weak*, and vice versa.

Note that these ‘external’ (between world states) and ‘internal’ (between agent states) transitions occur independently, so the dynamics can be written in terms of a Kronecker product between the external and internal dynamics, *𝒫*(*S*_*n*+1_|*S*_*n*_, *a*_*n*_) = *𝒫*(*X*_*n*+1_|*X*_*n*_) ⊗ *𝒫*(*Y*_*n*+1_|*a*_*n*_) (Fig. 1C).

Some exceptions to the preceding apply. Firstly, the transition between *X*_*N*−1_ and *X*_*N*_ is such that any belief in the *on* state at the end of time step *N* − 1 is transferred to *post* at the start of *N* (Fig. 1D). Secondly, it is certain that each trial begins in *pre*, but we assume that on transitioning from the *N* th step of trial *t* to the 1st step of trial *t* + 1, there is some probability *δ* of reverting from *strong* to *weak* attentional state (Fig. 1E). One might think of this as allowing the possibility of a passive or spontaneous “decay” process. We generally set this decay probability to be small (*δ* = 0.001).

#### 2.1.4 Observations

Depending on the sequence of states — and also, crucially, on the agent’s actions — the agent receives a sequence of observations *O*_1_, *O*_2_, …, *O*_*N*−1_. As mentioned above, we assume that no observation is provided at *N*, where the agent must report whether there was a signal or not during the trial. Observations carry information about whether there is currently a signal or not, but just how informative they are is determined by choice of attentional action. In particular, if the agent chooses to attend *weakly* on the current time step, then we assume:

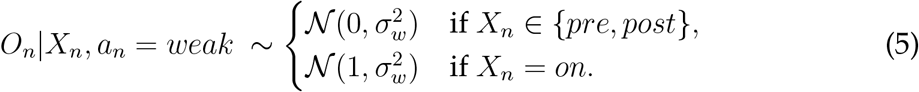

If the agent chooses to attend *strongly*, then we have

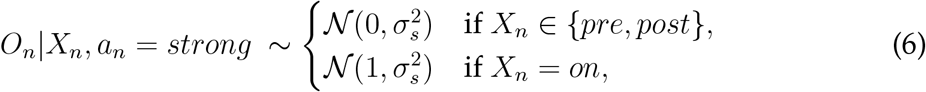

where 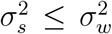. In other words, observations are always drawn from a normal distribution, with a mean of either 0 (if no signal) or 1 (if signal), and these observations are less ‘noisy’ in the *strong* than in the *weak* attentional state (Fig. 1B, right). Note that variability is assumed not to depend on whether the signal is present or not.

#### 2.1.5 Costs and rewards

While *strong* attention yields better quality information, we assume this comes at a cost, which may additionally depend on the current attentional state. Costs are always treated as non-positive (≤ 0) reward values. The only positive reward available in the task comes from correctly reporting the presence or absence of a signal.

We denote the reward function *R*_*n*_(*X, Y, a*), and assume that this itself is a sum of two terms,

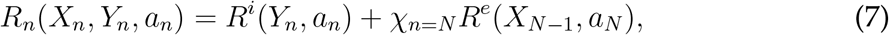

where *R*^*i*^(*Y*_*n*_, *a*_*n*_) gives the ‘internal’, attentional cost associated with attentional action *a*_*n*_ in attentional state *Y*_*n*_,

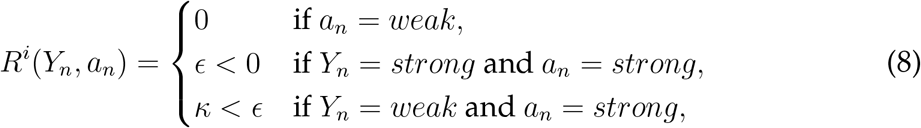

and *R*^*e*^ gives the ‘external’ reward arising from whether it was correctly or incorrectly reported on step *N* (signified by *χ*_*n*=*N*_) that a signal was present or absent during the trial, which we simply define as

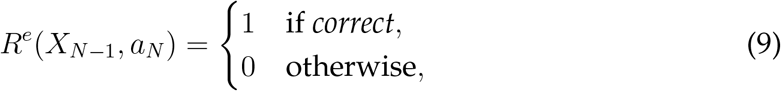

where the rectitude of a *no signal* or *signal* action *a*_*N*_ depends on whether *X*_*N*−1_ is *pre* or {*on, post*}. Note that the cost of attention only depends, in addition to attentional action *a*_*n*_, on the subcomponent *Y*_*n*_ (i.e., attentional state) of *S*_*n*_. It can be interpreted as follows: the cost of *weak* attention (*a*_*n*_ = *weak*) is zero, regardless of the previous action, while there is always a cost of *strong* attention (*a*_*n*_ = *strong*); furthermore, it is assumed that choosing to attend strongly given that one is already in the *strong* attentional state (*Y*_*n*_ = *strong*) is *less* costly than choosing to attend strongly given one is currently in the *weak* attentional state (*Y*_*n*_ = *weak*). The intuition behind this latter assumption is that such repeated action can also be conceptualized as maintaining a (non-default) internal state, which may incur less cost than switching between states (and in particular, from ‘default’/disengaged to ‘non-default’/engaged states; see Discussion).

#### 2.1.6 Belief MDP

In the current case, the agent’s posterior distribution over states given its observations, or ‘belief state’, coupled with memory of its most recent action, is sufficient to optimize behaviour. In particular, we can formulate the corresponding ‘belief MDP’, and use familiar dynamic programming methods to solve for the optimal policy.

Using simplified notation similar to Kaelbling et al. (1998), we denote by **b**_*n*+1_(*x*) the agent’s belief that *X*_*n*+1_ = *x* after seeing all the information *O*_1:*n*_ on time steps 1, …, *n*, including knowledge of the attentional actions *a*_1:*n*_ on those steps (and after one further probabilistic transition). (Strictly, **b**_*n*+1_(*x*), is a deterministic function of *O*_1:*n*_, *a*_1:*n*_ but, for notational convenience, we treat it as a random variable and do not write down this dependency.) That is, **b**_*n*+1_(*x*) := *𝒫*(*X*_*n*+1_ = *x*|*O*_1:*n*_, *a*_1:*n*_) = *𝒫*(*X*_*n*+1_ = *x*|*O*_*n*_, *a*_*n*_, **b**_*n*_). We then combine this belief over *X*_*n*+1_ with the known value of *Y*_*n*+1_ to make a state 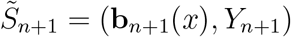 for what is called the belief MDP.

The transition and reward functions for the belief MDP are derived in the usual way. Specifically, the posterior probability over *X*_*n*+1_ given action *a*_*n*_ and particular observation *O*_*n*_ = *o*_*n*_ can be derived recursively via Bayes’ rule,

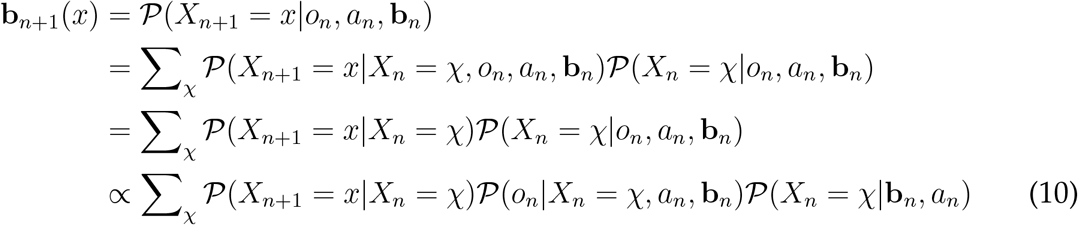

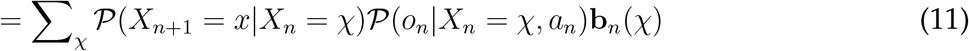

Following Kaelbling et al. (1998), we can think of this process in terms of the application of a state estimator function SE_*n*_(**b**_*n*_, *a*_*n*_, *o*_*n*_), which takes the initial belief state at *n*, along with the selected action and ensuing observation, and returns the resulting belief state. The transition function over belief states is then:

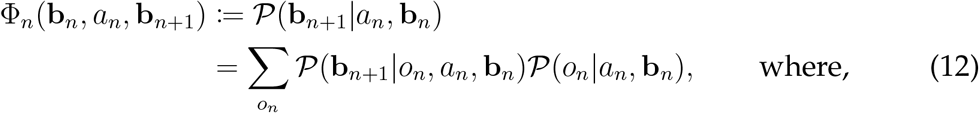

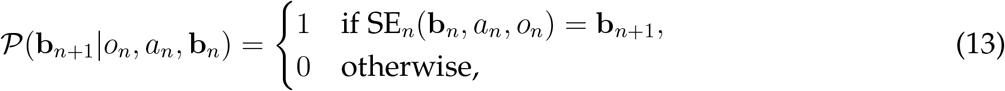

and where *𝒫*(*o*_*n*_|*a*_*n*_, **b**_*n*_) is the normalizing constant arising in Equation (10). This transition function, which marginalizes out the possible observations, is important when planning ahead, since the actual observations will not be known at that point.

Using the full state 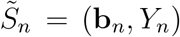, the reward function in the belief MDP is derived from the original reward function via:

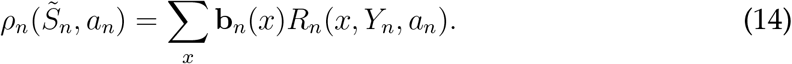

#### 2.1.7 Optimal behaviour

We consider optimal behaviour in two cases: the case of a single trial, and the case of the average reward rate over continuing trials.

For the single-trial case, a suitable objective is to determine a policy that maximizes the expected (undiscounted) return over the trial. Such an optimal policy can be found by simple backward induction. First consider the final time step *N*. Here, there is no observation to provide additional information and, furthermore, since there are no future trials to be considered, there is nothing to be gained by choosing other than the attentional action *a*_*n*_ = *weak* for any state. Then there is the additional choice of whether to report whether a signal was present or absent. The optimal decision here is simply to choose according to whether one’s belief about being in the presignal state is greater than 0.5 (report ‘no signal’) or not (report ‘signal’). From these considerations about the optimal policy at *N*, we can find the corresponding optimal values 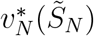, i.e., the maximum expected return for each possible state 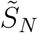 using (14), since the expected return is equal to the expected reward at this final state. Working backwards, we can then find the optimal values, and associated optimal actions, for all preceding time steps *n < N* using the Bellman optimality equation

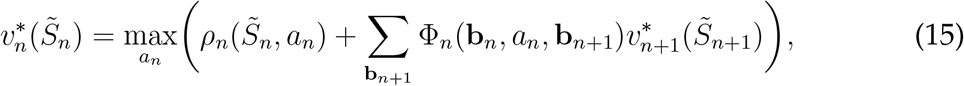

where 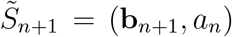. We thereby find a deterministic optimal policy 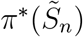, for all states 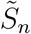, mapping from belief and attentional state *Y*_*n*_ at step *n* to an attentional action *a*_*n*_.

The multi-trial case is made slightly more complex by the need to consider future trials in addition to the current trial. This is because the actions in trial *t* can affect the starting state in trial *t* + 1. One suitable objective in this continuing case is to maximize the average rate of reward (Mahadevan, 1996), and we can solve for this maximum average reward rate *g*^*^ using value iteration to solve the system of Bellman optimality equations

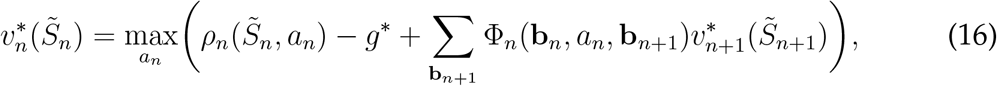

where it should be noted that 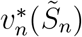 now refers to a *differential* value, based on the average-adjusted sum of (undiscounted) future rewards from each state (see, e.g., Sutton & Barto, 1998 for details).

In both single- and multi-trial cases, we approximate the continuous component of the belief state with a finite number of discrete states, specified by applying a regular grid with non-overlapping tiles of side Δ*b* (Lovejoy, 1991). Interpolation can then be used to extract values and actions for any other point in the belief space.

### 2.2 Attention and ACh: bridging hypothesis

When considering optimal (Bayesian decision-theoretic) behaviour, the representation and updating of uncertainty (as well as utilities) are obviously of key importance. It has been previously suggested (Dayan, 2012b; Dayan & Yu, 2006; Yu & Dayan, 2004, 2005) that the neuromodulators acetylcholine (ACh) and norepinephrine (NE) play an important, if circumscribed, role in representing (though not themselves calculating) expected uncertainty (i.e., variability in the world that is predicted) and unexpected uncertainty (i.e., variability that is not predicted) at both relatively slow (for learning) and fast (for inference) timescales. This complements other forms of representation, such as in population codes (Orbán, Berkes, Fiser, & Lengyel, 2016; Pouget, Beck, Ma, & Latham, 2013). Our interest here is primarily in fast ACh signalling, and its possible roles in inference (Hasselmo & Sarter, 2011; Sarter & Lustig, 2019, 2020).

The earlier ideas about the role of ACh in inference were that it directly reports cases in which prior expectations about the world are vague (for example, in the SAT, arising from variable timing and the low SNR of sensory inputs). In such cases, it is appropriate to weight external information more heavily (i.e., boosting bottom-up over top-down information) and potentially to improve one’s prior from this new information. However, that work involved Bayesian inference rather than Bayesian decision theory, and so did not consider the additional possibility of an attentional action — i.e., the possibility that the SNR of the input could advantageously, though possibly expensively, be changed to increase the net reward rate. We suggest that ACh also plays this additional role in the SAT, such that the release of ACh can be treated as an internal action rather than as a passive reporter of expected uncertainty. In some cases, these roles may be functionally indistinguishable.

We generally consider ACh to report *a*_*n*_ = *strong*. However, when we consider experiments in which ACh is manipulated directly by optogenetics (Gritton et al., 2016), we discuss some of the complexities of breaking the relationship between the optimally intended and the actual release of ACh — since there could then be mismatch between the assumed and actual qualities of the information that is being reported and integrated (see Section 3.2).

It is important to note that ACh is substantially heterogeneous (Gielow & Zaborszky, 2017; Záborszky et al., 2018). In the experiments we report, there are differential effects associated with sensory cortical and prefrontal cortical ACh. At present, it is not possible to be completely precise in the model about the release and effects of ACh — themes that we return to in the Discussion.

### 2.3 Plots and Measures

To facilitate understanding of subsequent results, we briefly introduce the main types of plot we use to understand model behaviour. Key measures of performance are also described.

One type of plot concerns which attentional action is optimal in each possible belief state **b**_*n*_. Since at any time step *n* we have 3 possible trial states, *X*_*n*_ ∈ {*pre, on, post*}, and a belief state must sum to 1, then the belief space at any time can be visualized as a 2-dimensional simplex (i.e., a triangle). The lower panels of Figures 3A–C, below, depict the belief space at a particular time step, and show the value (Fig. 3A, lower) and the optimal attentional actions (Figs 3B–C, lower) for each belief state, conditioned on the current value of *Y*. Each time step in a trial has a belief space of this form, and so we can visualize changes in value or policy over the course of a trial by appropriate concatenation of such plots (as in the upper panels of Figs 3A–C).

**Figure 3:**
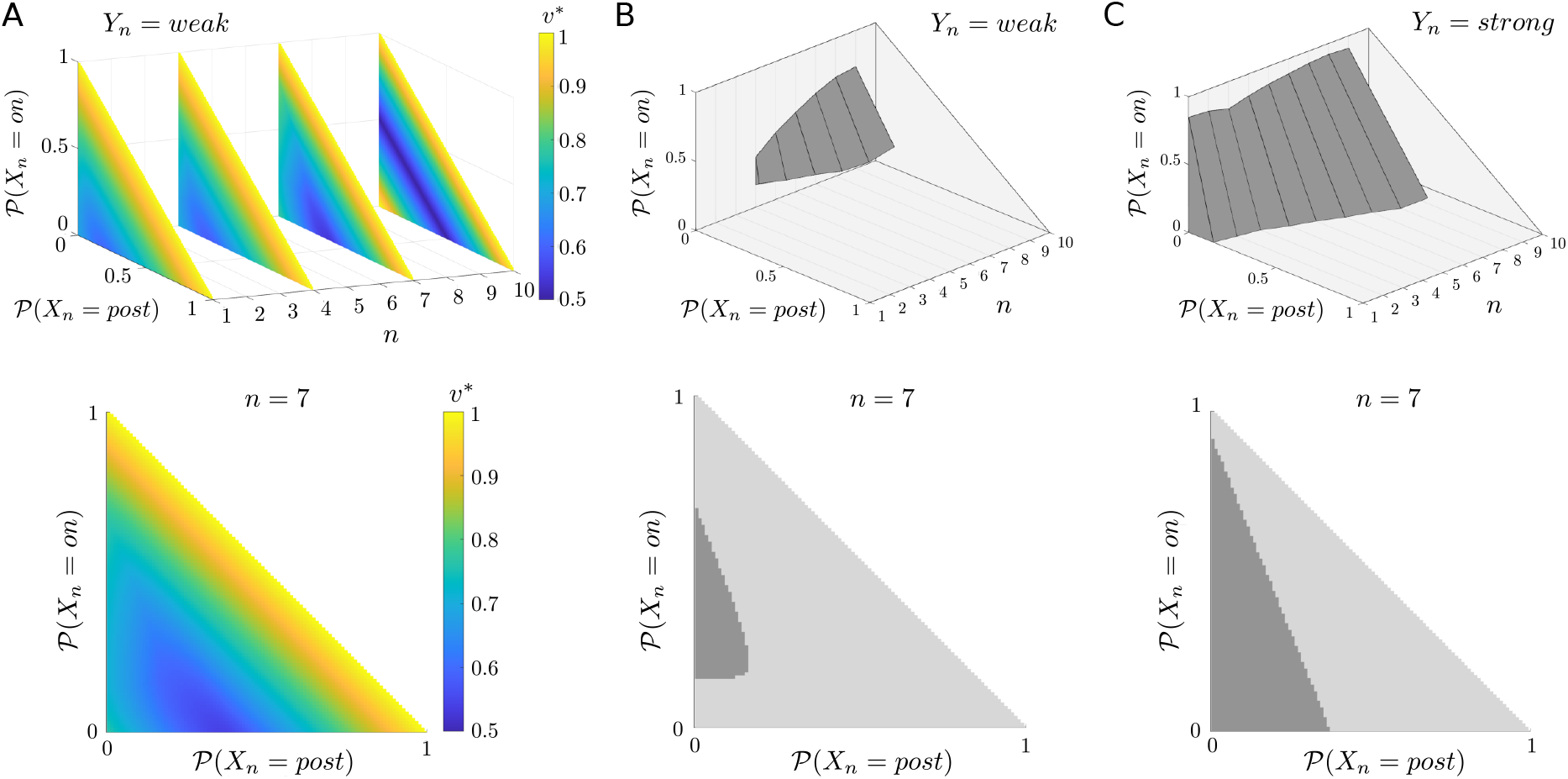
Optimal values and actions for the single-trial, frequent-signal regime. (A) Optimal state values (*v*^*^) for a subsample of time steps (upper), and expanded for the particular case of *n* = 7 (lower); in this case, values are always conditioned on current occupation of the *weak* state, *Y*_*n*_ = *weak*. At the decision state (*n* = 10), the value of being more certain about whether there has been a signal or not is clear, since these belief states have higher *v*^*^. This value is the same regardless of whether one is certain that there was or was not a signal. For earlier time steps, the highest values occur only where one is certain that a signal is, or has been, present. (B–C) The associated optimal policies (upper plots) conditioned on currently occupying (B) the *weak* attentional state (*Y*_*n*_ = *weak*) or (C) the *strong* attentional state (*Y*_*n*_ = *strong*); lower plots for time step *n* = 7. Darker regions bound the set of beliefs for which choosing *strong* is optimal; in general, this region grows over time.

Another plot type shows the average evolution of the belief state over a trial, along with the average evolution of the attentional state. Figure 4A (left) shows one such case: note that at any time, the sum of beliefs in each possible state (*pre, on, post*) sums to 1 (the upper plot); the evolving attentional state — we simply label the *y*-axis ‘attention’ — is more precisely the probability of selecting *strong* at each time step, *𝒫*(*a*_*n*_ = *strong*) (lower plot). The probability of choosing *strong* at each step is alternatively rendered in terms of shaded squares when systematically varying parameters (e.g., Fig. 8A, upper left), along with the ‘marginal’ expected number of steps per trial in which *strong* is chosen (e.g., Fig. 8A, lower left).

**Figure 4:**
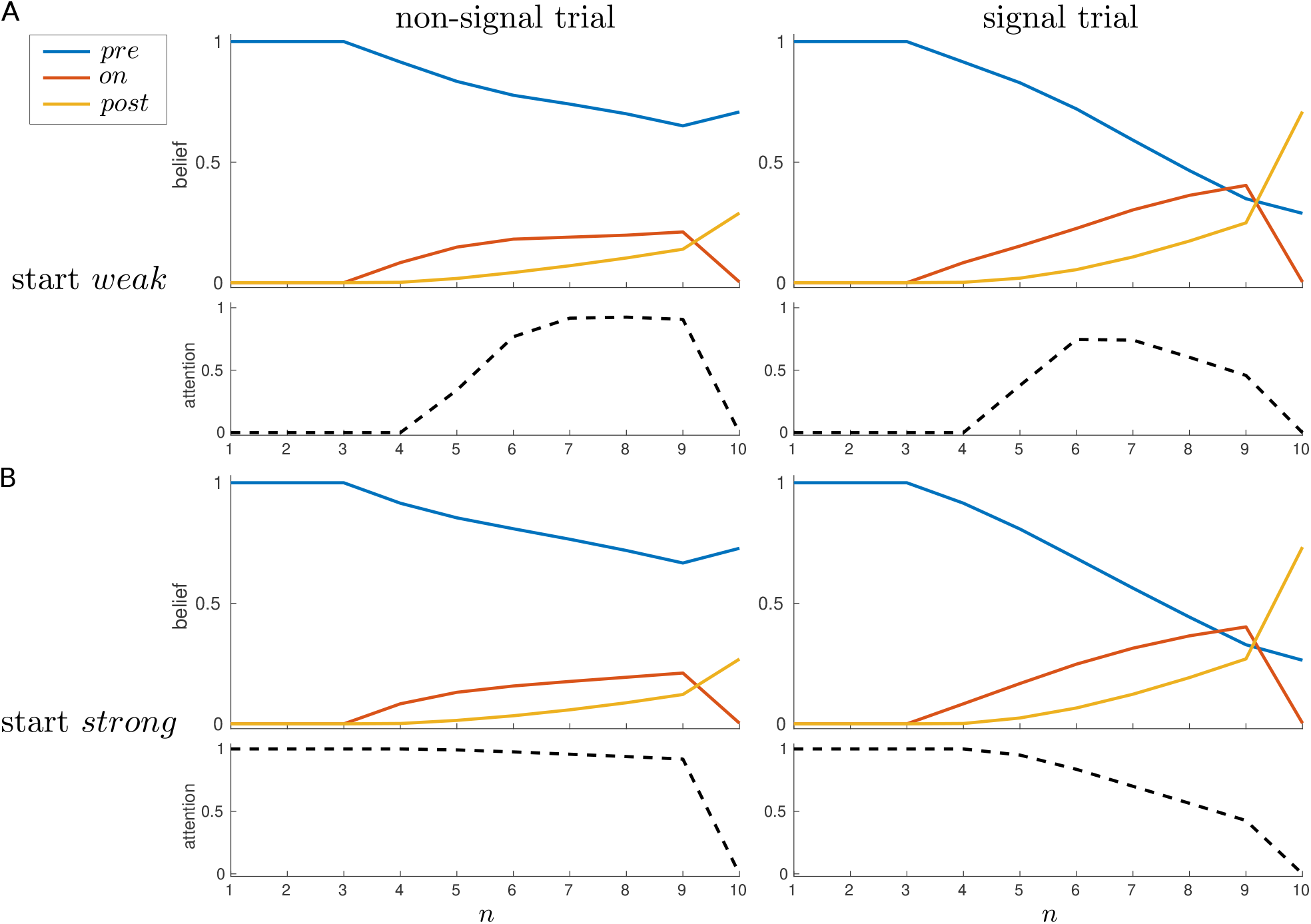
Dynamics of belief and attention, single-trial case. Average beliefs and attention (i.e., *𝒫*(*a*_*n*_ = *strong*)) for a single trial, starting in either (A) the *weak* or (B) the *strong* attentional state, separated by whether the trial contains a signal (right panels) or not (left panels).

**Figure 5:**
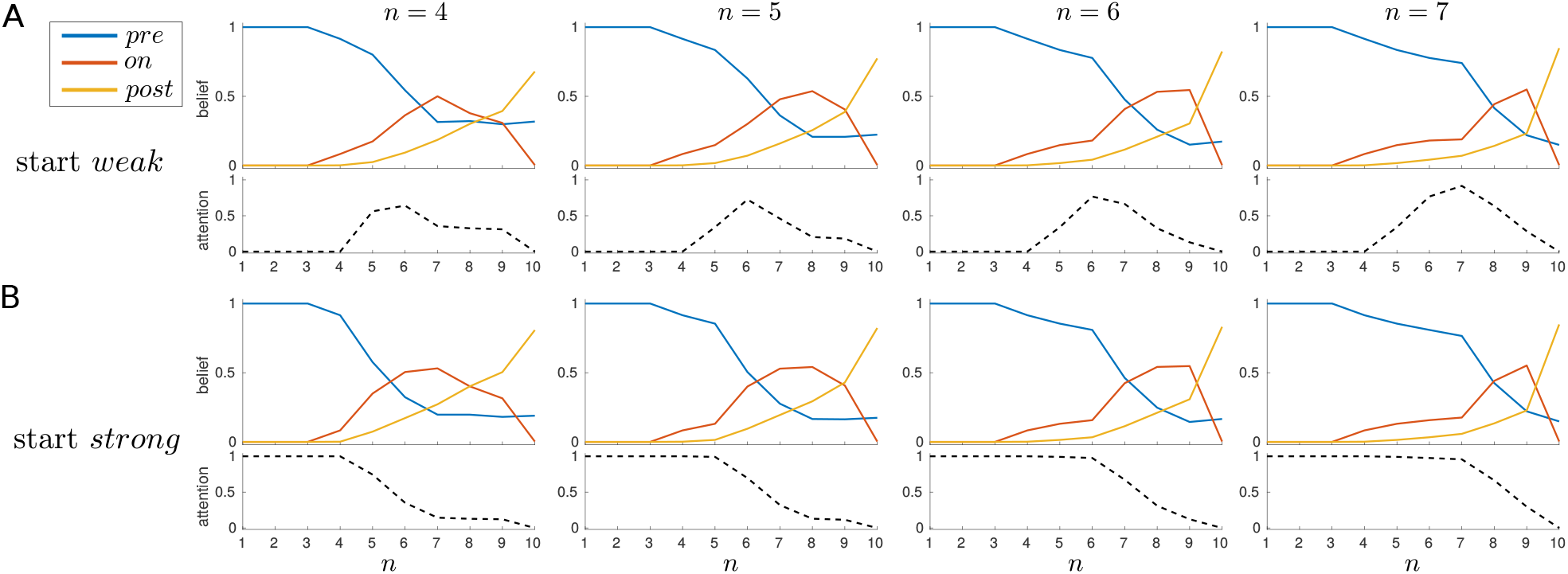
Modulation of dynamics by signal onset, single-trial case. Average beliefs and attention for signal trials in which the signal always has a duration of 3 time steps, but with different onsets, as indicated at top. (A) Start *weak*; (B) start *strong*.

**Figure 6:**
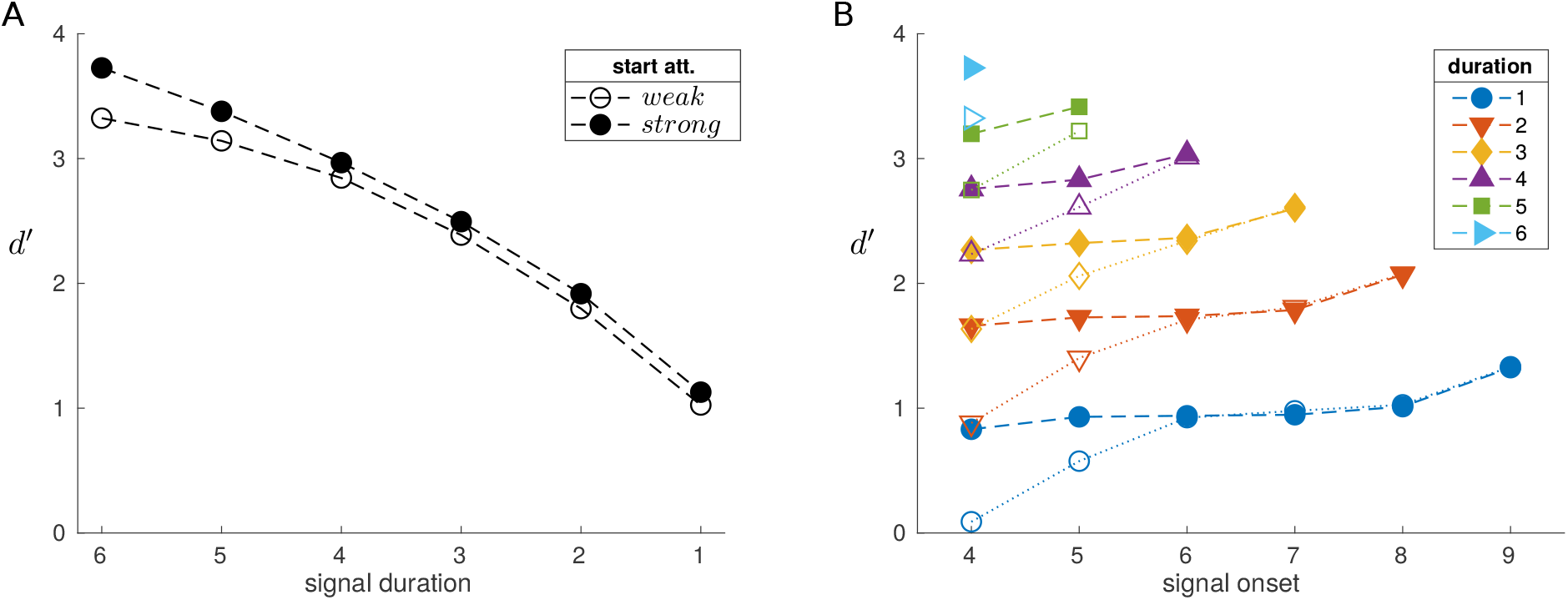
Sensitivity is higher for longer, later signals. (A) Sensitivity *d*^*′*^ is higher for longer signals; starting the trial in the *strong* attentional state leads to enhanced sensitivity. (B) *d*^*′*^ is also modulated by signal onset; a signal of a given duration with a later onset is better detected (filled markers when starting the trial in the *strong* state).

**Figure 7:**
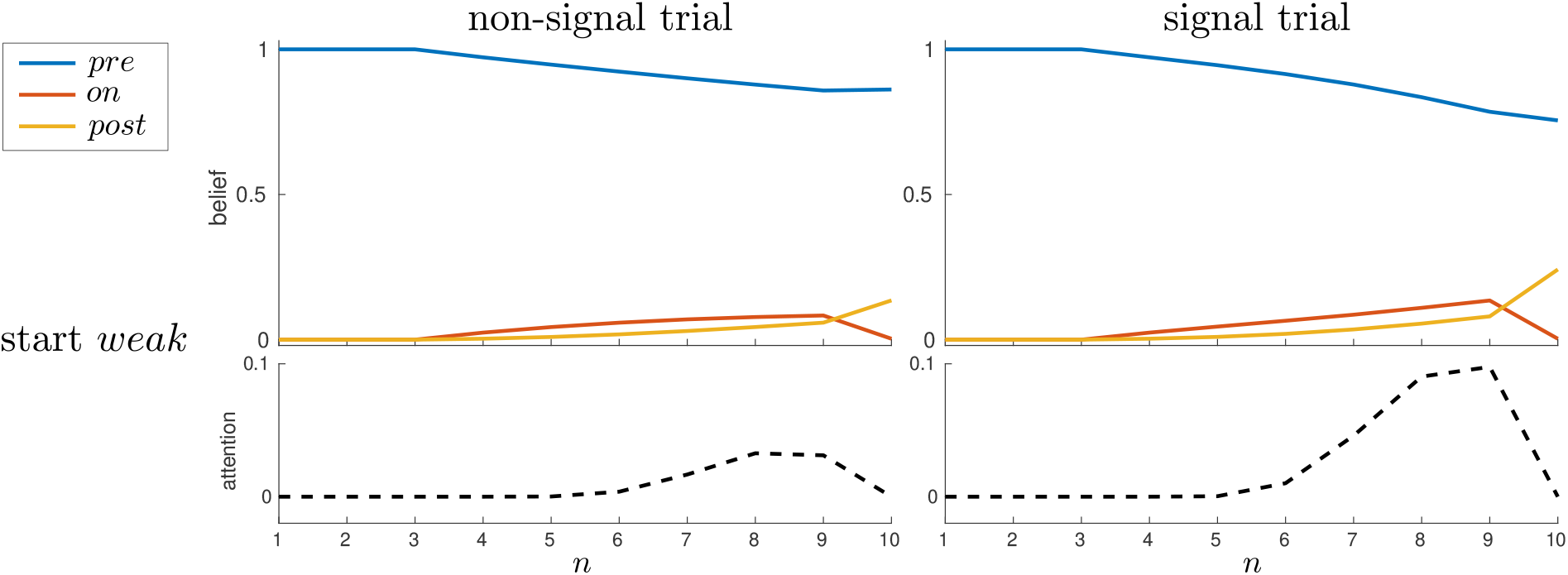
Greater engagement of *strong* attention on signal trials when signals are rare. Average beliefs and attention for a single trial, started in the *weak* attentional state, depending on whether a signal is absent (left panels) or present (right panels). The prior probability of a signal is low, *p*_1_ = 0.15; *ϵ* = 0.005. Parameters otherwise as in previous example.

**Figure 8:**
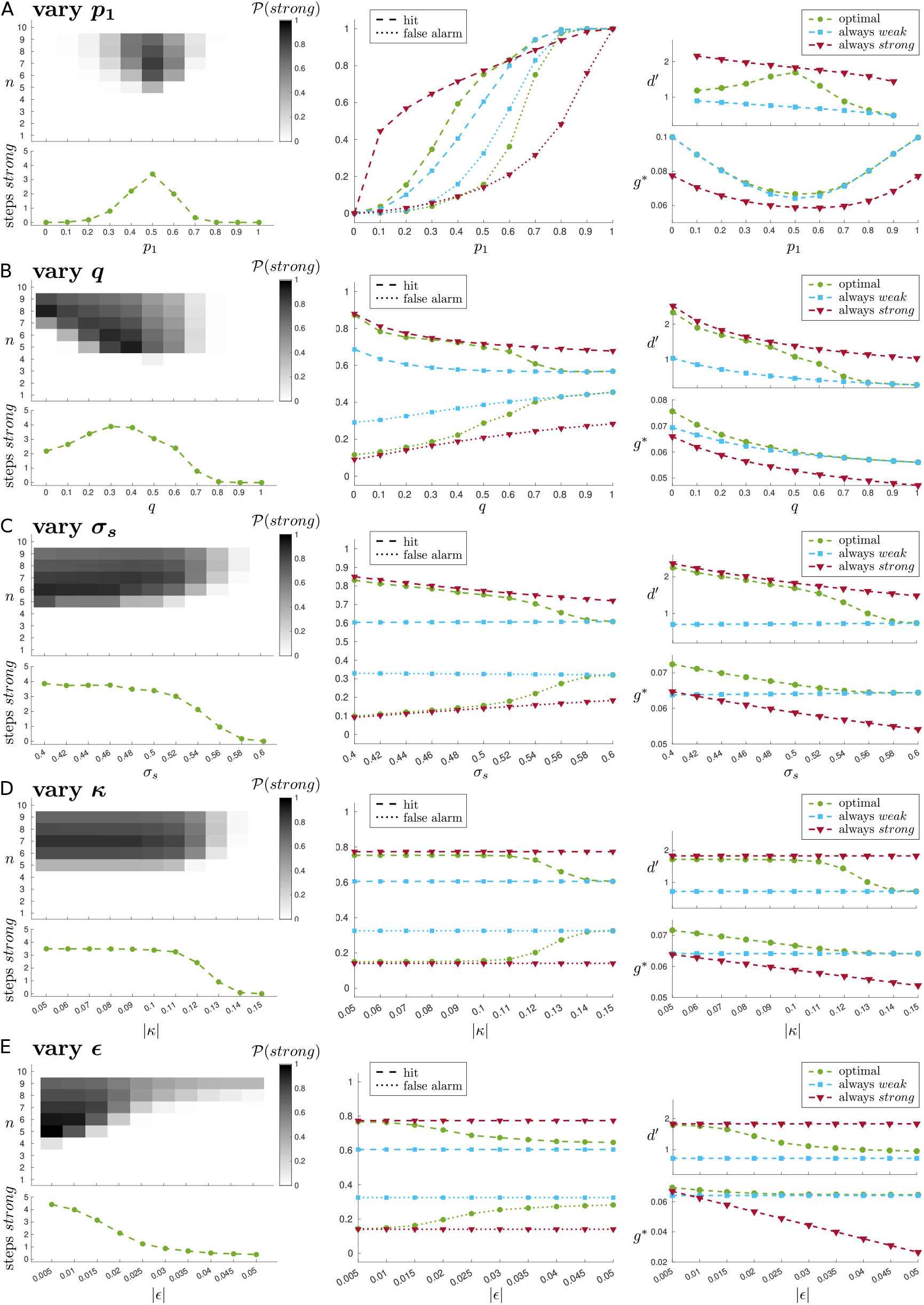
Effects of varying model parameters: (A) signal probability, *p*_1_; (B) signal offset parameter, *q*; (C) quality of information in *strong* attentional state, *σ*_*s*_; (D) absolute cost of *weak* → *strong* shift, |*κ*| ; (E) absolute cost of maintaining *strong* attention, |*ϵ*|. For each parameter, we show the expected occupation of the *strong* state (left; upper plot shows probability of occupying *strong* at each time step, while lower plot shows the ‘marginal’ expected number of steps in *strong* state); the hit rates and false alarm rates (middle); and the sensitivities *d*^*′*^ derived from the hits and false alarms, and average reward rates *g*^*^ (right). For the middle and right plots, we compare the performance of the optimal policy with the ‘always-*weak*’ and ‘always-*strong*’ policies (see main text). Parameters varied one at a time, with other parameter values fixed at ‘default’ values: *p*_1_ = 0.5; *q* = 0.2; *σ*_*w*_ = 1; *σ*_*s*_ = 0.5; *κ* = − 0.1; *ϵ* = − 0.014; *N*_0_ = 3; *N*_1_ = 6.

Predicted ACh activity under the model (Figs 12A–B, right panels) is based on the assumption that ACh reports *a*_*n*_ = *strong* (see Section 2.2 above), and so is similarly derived directly from *𝒫*(*a*_*n*_ = *strong*). The only additional subtlety is that the ACh transients reported by Howe et al. (2013) (Figs 12A–B, left panels) are measured relative to a pre-signal baseline (2 s before signal onset, or analogous period on a non-signal trial), and we therefore similarly report predicted changes in ACh relative to a baseline obtained by averaging attentional state over 3 time steps before the signal onset (signal trials) or the average signal onset time (non-signal trials).

**Figure 9:**
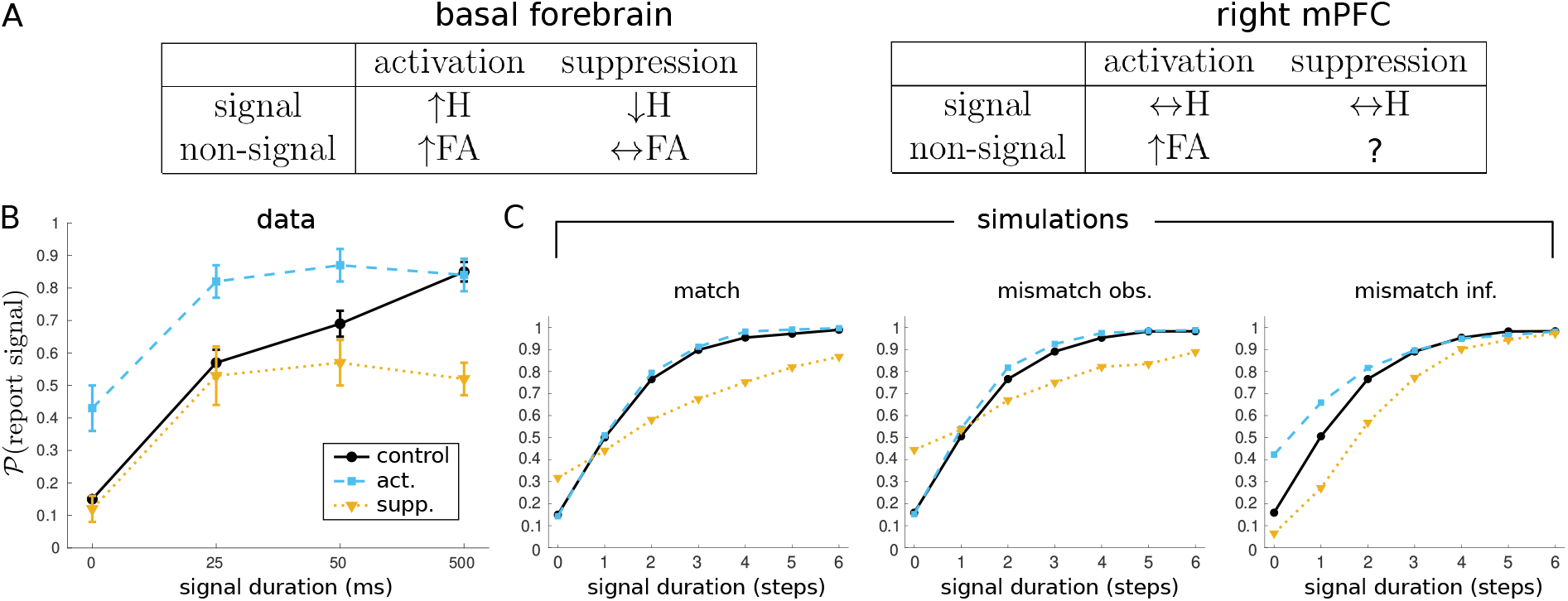
Effects on hits and false alarms of optogenetic activation and suppression. (A) Summary of effects of optogenetic manipulations on hits (H) and false alarms (FA) in mice performing the sustained attention task (SAT; replotted from Gritton et al., 2016). (B) Effect of activating or suppressing basal forebrain at maximum power (replotted from Gritton et al. (2016)). (C) Effects of different ‘optogenetic’ manipulations in the model using the default set of parameters; each data point reflects 10,000 simulated trials. In each plot, ‘control’ performance is always the same, and reflects the case where observations are generated under the optimal attentional policy, and also believed to be so generated for the purposes of inference. Left (‘match’): actual and believed properties of observations matched, but generated either under an always-*strong* policy (‘act.’), or an always-*weak* policy (‘supp.’). Middle (‘mismatch obs.’): observations always believed to be generated under the optimal attentional policy, but actually generated under an always-*strong* (‘act.’) or always-*weak* (‘supp.’) policy. Right (‘mismatch inf.’): observations actually generated under the optimal attentional policy, but believed to be generated under an always-*strong* (‘act.’) or ‘always-*weak*’ (‘supp.’) policy.

**Figure 10:**
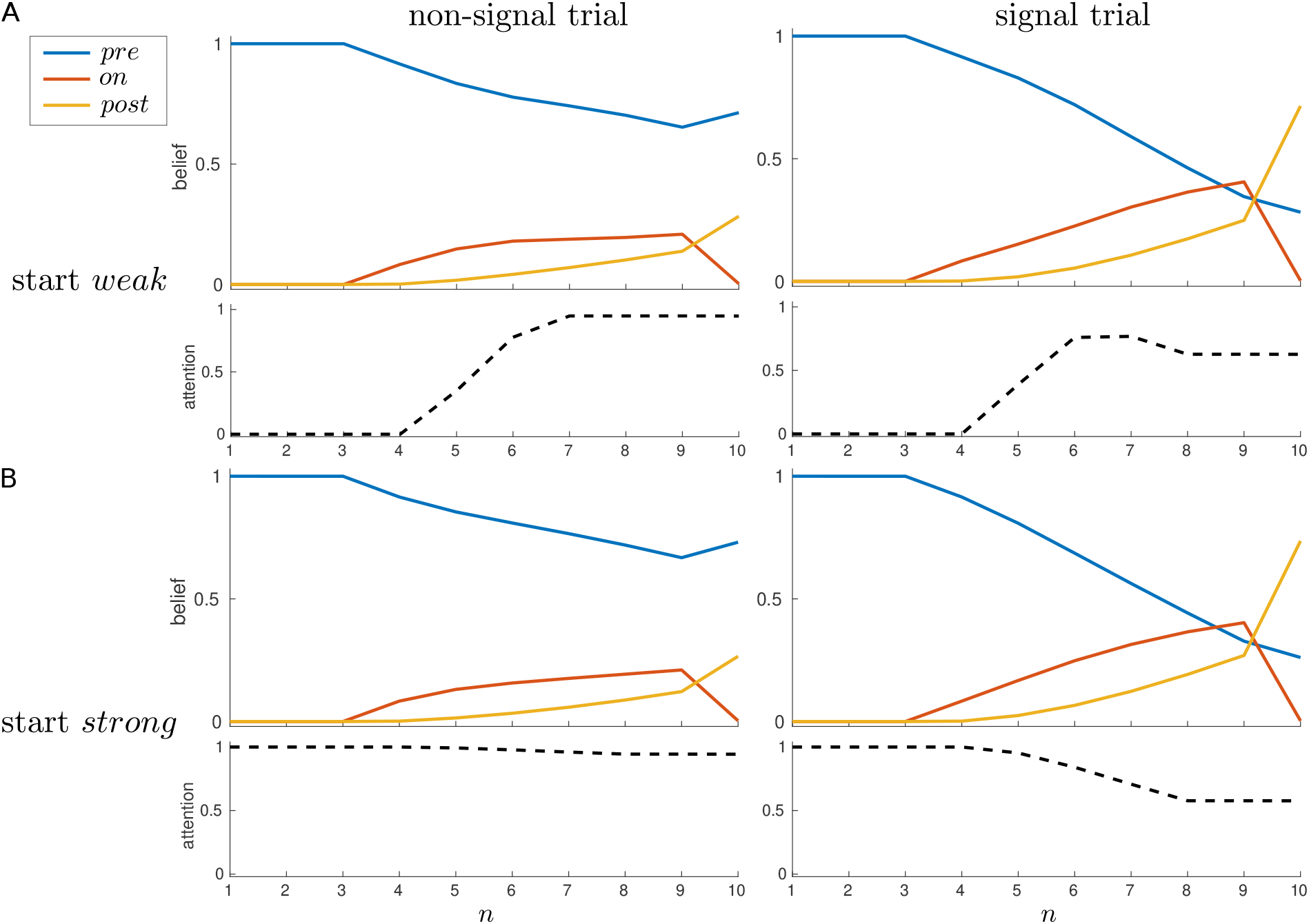
Maintenance of *strong* attention in the multi-trial case. Average beliefs and attention for the multi-trial case, starting in either (A) the *weak* or (B) the *strong* attentional state, separated for non-signal (left panels) and signal (right panels) trials.

**Figure 11:**
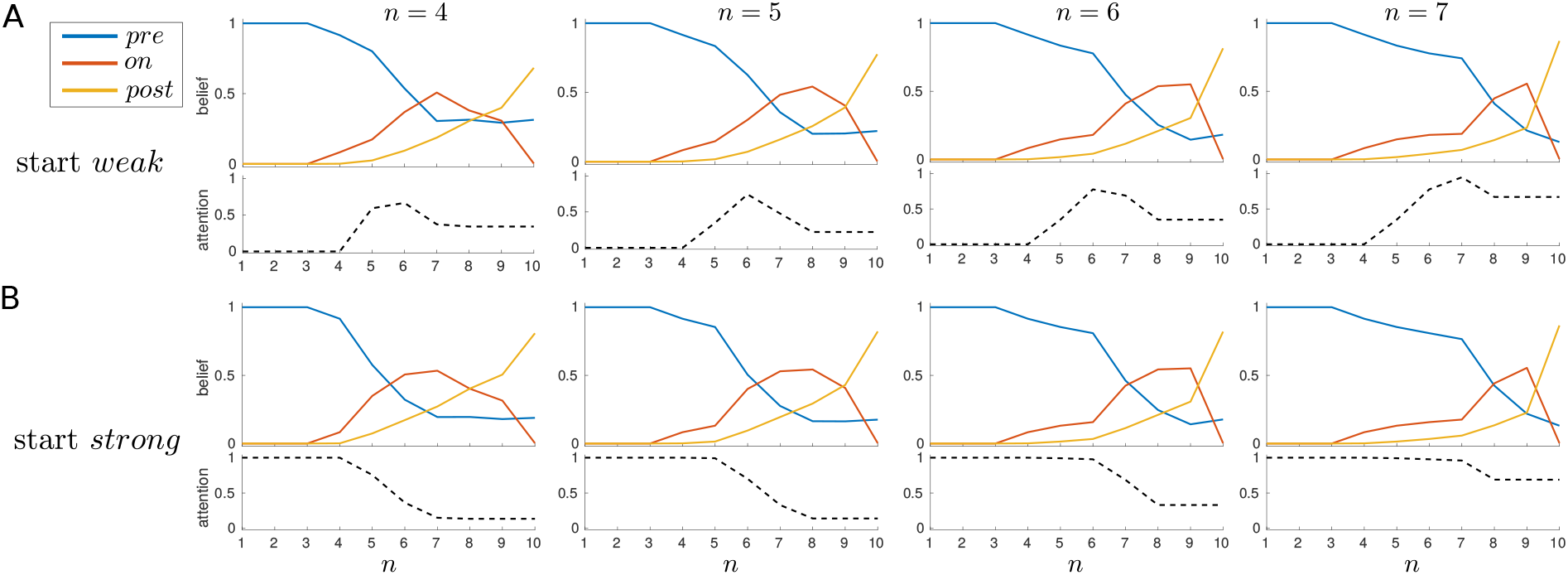
Maintenance of *strong* attention is modulated by signal onset. Average beliefs and attention for a trial with a signal of duration 3 time steps but with different onsets, as indicated at top. (A) Start in the *weak* attentional state; (B) start in the *strong* attentional state.

**Figure 12:**
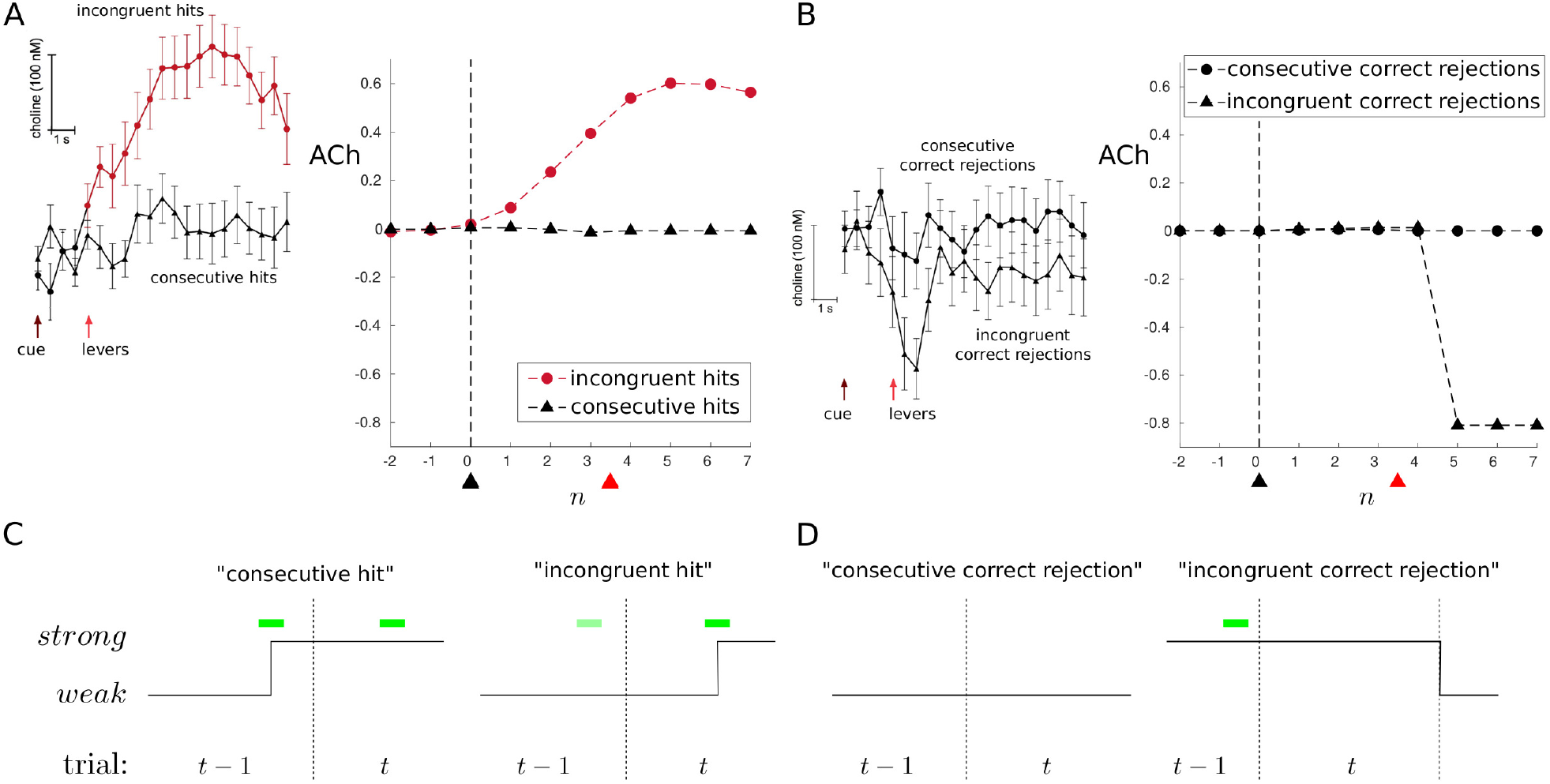
ACh signals measured in rat mPFC by Howe et al. (2013) during the sustained attention task (SAT). (A) Signal trials. Left: a significant increase in ACh on signal trials in which an animal correctly reported a signal (i.e., hit trial), but only when a non-signal was reported on the previous trial (‘incongruent hit’) and not if the previous trial was also a hit trial (‘consecutive hit’). From Howe et al. (2013). Right: simulation results. (B) Non-signal trials. Left: no increase in ACh on correct rejection trials; a non-significant trend for a transient decline in ACh on a correct rejection trial if preceded by a hit trial (‘incongruent correct rejection’). From Howe et al. (2013). Right: simulation results. In the experiments, ACh signals are relative to a presignal baseline (period of 2s before signal onset, or analogous period on non-signal trial), and aligned to signal onset or analogous time point for non-signal trials (black arrows); average time of levers’ insertion following cue (red arrows). In the simulations, ‘ACh’ is measured relative to a baseline obtained by averaging attentional state over 3 time steps before the signal onset (signal trials) or the average signal onset time (non-signal trials). Vertical dashed line and black arrow indicate cue; red arrow indicates average time of trial end. Averages reflect 10,000 simulated trials. Prior signal probability *p*_1_ switches between 0.1 and 0.15, with parameters otherwise fixed as follows: *q* = 0.2, *σ*_*w*_ = 1, *σ*_*s*_ = 0.5, *κ* = −0.1, *ϵ* = −0.005, *δ* = 0.001. (C) Left: if a signal has prompted a *weak* → *strong* attentional shift (and detection) on trial *t*−1, this *strong* attention may persist into trial *t*, where detection is associated with no change in attentional state. Right: if the *weak* attentional state is currently occupied, a *weak*→*strong* shift is most likely to occur on a (detected) signal trial; *t* − 1 may have no signal or a signal that is undetected (i.e., a miss trial). (D) Left: consecutive correct reports of no signal are not expected to be associated with attentional change. Right: if a signal (and detection) means that the *strong* attentional state is occupied at the end of trial *t* − 1, it may persist into trial *t*; however, if there is no signal in this next trial, then there may be a reversion back to the *weak* attentional state (i.e., a relative decrease).

In addition to these plots, we report the common sensitivity measure *d*^*′*^ used in signal detection theory (Green & Swets, 1966; MacMillan & Creelman, 2005),

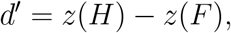

where *H* is the hit rate (i.e., the probability of reporting a signal given a signal was presented), *F* is the false alarm rate (i.e., the probability of reporting a signal given no signal was presented), and *z*(·) is the quantile function of the standard normal distribution. Note that we can make such measures conditional on the onset and duration of a signal (as in Fig. 6); that is, while *F* is the probability of reporting signal when none has been presented, we can separately consider the hit rates *H* for signals of different duration, disregarding onset (Fig. 6A), or even examine *H* for all possible onset-duration combinations (Fig. 6B). In Figure 8, we report hit and false alarm rates (middle plots) in addition to *d*^*′*^ (upper right plots).

## 3 Results

We consider both internal and external decision-making over the two key timescales in the task. The most obvious timescale is that associated with each individual trial. Within a trial, the agent has to accumulate evidence about the presence of the signal, decide whether and when to engage or disengage attention, and finally report their external choice (report ‘signal’ or ‘no signal’). However, we are also concerned with what occurs *across* trials. Although from a signal detection viewpoint each trial is independent, and so the actual presence of a signal on one trial bears no relationship to its presence on previous or future trials, the agent’s attentional state is allowed to persist across trials. Indeed, as we noted in the Introduction, it might be substantially more expensive to engage *strong* attention than to maintain it. Then, if attention is already engaged on trial *t*, it might be optimal to pay the cost of sticking in the *strong* state for trial *t* + 1 and thereby avoid the even greater cost of having to re-engage this state, if necessary. In turn, this would mean that the choice to engage attention on trial *t* can be partly justified by the benefits of *strong* attention on trial *t* + 1. Thus, we need to consider an inter-trial timescale as well as an intra-trial one. We start with the case of an isolated trial, and then treat cross-trial influences.

### 3.1 Single trial

When does it make sense to engage (*weak* → *strong*) or disengage (*strong* → *weak*) attention within a trial, if we only consider optimizing behaviour in a single trial (i.e., ignoring the effect of choice on future trials)? We expect multiple model parameters to interact to determine the answer, since the probability that a signal is on and its temporal extent (determined by *p*_1_, *q* and *N*_1_), the relative qualities of information available in different attentional states (*σ*_*w*_, *σ*_*s*_), and the costs of *strong* attention (*κ, ϵ*) can all be expected to play a role. To gain some insight, we examine a particular regime in detail (these will be our ‘default’ parameters) before more systematically examining the effects of varying parameters.

#### 3.1.1 Example: frequent signals

In our first, ‘default’ regime, we assume that a signal is as likely as not, *p*_1_ = 0.5, with *q* = 0.2. In terms of the relative quality of observations, we set *σ*_*w*_ = 1 and *σ*_*s*_ = 0.5. For costs, we assume the cost of a *weak* → *strong* shift to be moderately costly, *κ* = −0.1, and the cost of maintaining *strong* attention to be substantially cheaper, *ϵ* = −0.014. Finally, for convenience, a trial is relatively short, with *N*_0_ = 3 and *N*_1_ = 6 (so that, in total, a trial includes *N* = 10 time steps).

The upper plot in Figure 3A shows the optimal values of each belief state at selected time steps (1, 4, 7 and 10) when occupying the *weak* attentional state; the lower plot highlights step *n* = 7 for clarity. At the decision state (*n* = 10), state values are higher for more certain beliefs (where one is more likely to be correct), and lowest where there is most uncertainty (where one can expect to perform at chance). Moving backwards in time, the value remains high when convinced there is, or has been, a signal (i.e., when *𝒫*(*X*_*n*_ = *on*) + *𝒫*(*X*_*n*_ = *post*) ≈ 1). This is not true where there is strong belief in the pre-signal state (i.e., *𝒫*(*X*_*n*_ = *on*) + *𝒫*(*X*_*n*_ = *post*) ≈ 0) — since this latter conviction does not preclude the possibility of a signal arriving in the future.

Figures 3B and 3C show the associated optimal policies when occupying the *weak* and *strong* attentional states, respectively. In each case, the dark, bounded region depicts the set of states at each time step where it is optimal to choose *strong*. (There are only planes at each discrete time step; the interpolated rendering is for didactic clarity.) This region grows over time, in accordance with the growing probability of a signal being on. In neither case, however, is there any reason to select *strong* in the final decision state (*n* = *N* = 10) — this is because no information is provided at this state, and so it can have no bearing on performance in the current trial (and any impact on performance in future trials is not currently considered).

Again, if we consider working backwards through time, we see in the *weak* attentional state (Fig. 3B) that choosing *strong* is generally worthwhile if one is uncertain about whether a signal is present, and belief in the post-signal state is low; this trend extends backwards in time, for a shrinking set of beliefs, until at *n* = *N*_0_ = 3 there is no belief state where choosing *strong* pays off. As already mentioned, the increase with time in the set of beliefs for which choosing *strong* is optimal is because of the non-uniform probability of a signal being present at any given time step (cf. Fig. 2B): since *q <* 1, a signal typically stays on for more than 1 time step, and it can therefore be economic to choose *strong* only later rather than earlier.

Conversely, once in the *strong* attentional state, it can be optimal to choose to continue in this state (Fig. 3C). Again, on the penultimate time step, this occurs where there is uncertainty about being in the signal state, and belief in the post-signal state is low. Moving backwards in time, this region tends to shrink, and notably incorporates the region where belief in the pre-signal state is high (i.e., *𝒫*(*X*_*n*_ = *on*) + *𝒫*(*X*_*n*_ = *post*) ≈ 0) — again, since a current belief that one is in the pre-signal state is no guarantee that a signal may not occur in the future. Note that this includes the initial *N*_0_ = 3 time steps: even if in the current case we would never expect a trial to begin in the *strong* state, if this were to happen, then this state would be maintained over the initial *N*_0_ steps, despite the certainty that the pre-signal state is occupied. Where it is certainly *not* worthwhile to continue to choose *strong* is when sufficiently certain that one is in the signal or post-signal state.

While such plots of the optimal values and policy are instructive, they do not by themselves tell us how beliefs or attentional state evolve over time. For example, since each trial begins with certainty that one is in the pre-signal state, *𝒫*(*X*_1_ = *pre*) = 1, and this certainty extends over the *N*_0_ initial time steps, there is no change in belief during this time.

For an alternative view, Figure 4 shows the average belief trajectories and average probability of choosing *strong* attention throughout a trial, separated by whether a signal did or did not occur, and by whether the trial was started in a *weak* (*Y*_1_ = *weak*) or *strong* (*Y*_1_ = *strong*) attentional state (the difference between which is quite subtle for the current parameters). In all cases, the initial certainty about the pre-signal state decreases, and on average ends up above 0.5 for a non-signal trial, and below 0.5 for a signal trial.

When starting in the *weak* state (Fig. 4A), the probability of choosing *strong* increases at *n* = 5, whether (right) or not (left) there is actually a signal in the trial, and returns to zero at the decision state (*n* = 10). While initially the probability of choosing *strong* is similar for non-signal and signal trials, one can see that this probability reaches a higher level in the non-signal case, and is maintained up to the penultimate time step. By contrast, the probability of choosing *strong* on a signal trial decreases from time step 6 onwards. This difference comes from the asymmetry already mentioned: once a signal has been detected, there is no reason to pay the cost of remaining in the *strong* attentional state, since higher quality sensory information cannot change the agent’s mind — a signal can either stay on or switch off. However, if the signal has not yet been detected, it remains theoretically possible (until *n* = 10) for it to turn on. We also see this effect when starting in the *strong* state (Fig. 4B): whether a signal is absent (left) or present (right), *strong* is initially maintained, but the average probability of choosing *strong* decreases more rapidly on a signal trial.

Figure 5 further illustrates this effect by averaging signal trials in which the signal is always of duration 3, but comes on at different times (i.e., at *n* = 4, *n* = 5, *n* = 6, or *n* = 7). Regardless of whether the trial is started in the *weak* or *strong* attentional state, an earlier onset effectively means that a decision is made earlier that a signal is indeed present, and so attention can be relaxed sooner within the trial. On the other hand, being sure that there has been no signal so far is no guarantee that one will not arise in the future — and so, once in the *strong* attentional state, it can make sense to maintain this heightened state of vigilance.

As one would expect, discrimination performance is better for longer signals, and starting in the *strong* state leads to improved sensitivity (Fig. 6A). At a finer grain, sensitivity is also modulated by signal onset (Fig. 6B). In general, regardless of whether one starts the trial in a *weak* or *strong* attentional state, the later a signal arrives, the more likely it is to be detected. This can ultimately be attributed to the increasing hazard function of a signal over the course of a trial, which leads the optimal policy more frequently to have attention be *strong* during the signal. Note also that the improvement in sensitivity when starting in the *strong* attentional state is due to improved detection of signals that arrive early in the trial.

#### 3.1.2 Example: rare signals

In the preceding example, we set *p*_1_ = 0.5, meaning that in expectation, half of the trials will contain a signal. One observation was that a *weak* → *strong* shift was likely to occur whether a signal was present or not, and indeed, the probability of occupying the *strong* state on a given step was generally higher on a non-signal trial (cf. Fig. 4A). Figure 7 shows an example in which occupation of the *strong* state is instead higher for a signal trial. This regime, in which signals are expected to be rare (we set *p*_1_ = 0.15), will also be relevant when we consider sequential effects below (Section 3.3.2). Although the overall probabilities of occupying the *strong* attentional state are much lower (note the *y*-axis scale for attention), one sees that the engagement of *strong* attention is more likely on a signal than a non-signal trial. Note, however, the much greater probability of missing a signal, since, again in expectation, *𝒫*(*X*_*N*_ = *pre*) *>* 0.5 (leading to a report of ‘no signal’) on a signal trial.

#### 3.1.3 Effects of varying model parameters

We next measure the effects of varying the default set of parameters systematically, one at a time. In reporting hit/false alarm rates, sensitivities, and average reward rates, we compare the optimal policy to two extreme cases: the ‘always *weak*’ policy, which enforces constant occupation of the *weak* attentional state; and the ‘always *strong*’ policy, which means the *strong* attentional state is always occupied.

The overall results (Fig. 8) can be summarized as follows: for *p*_1_ and *q*, there are intermediate parameter values which lead to greatest engagement of *strong* attention; otherwise, as the relative improvement in quality of information from engaging *strong* decreases (i.e., as *σ*_*s*_ approaches *σ*_*w*_), or as the associated costs increase (i.e., |*κ*| and/or |*ϵ*| increase), then there is progressively less engagement of *strong* attention until a ‘breaking point’ is reached — when it is no longer paying the price of *strong* attention at all.

##### Signal probability, *p*_1_

When measuring the effect of varying signal probability *p*_1_ more systematically (Fig. 8A), we see that the highest occupancy of the *strong* attentional state occurs when there is most uncertainty about whether there will be a signal or not, i.e., *p*_1_ = 0.5 (Fig. 8A, left); this is also where *strong* attention is likely to be engaged earliest within a trial. As one would expect, the ability to engage *strong* attention generally leads to improved hit and false alarm rates (Fig. 8A, middle), i.e., improved sensitivity (Fig. 8A, upper right). Also as expected, *p*_1_ = 0.5 is where reward rate is lowest, but it is also where the ability to choose *strong* leads to the largest increase in reward rate compared the always-*weak* policy (Fig. 8A, lower right).

##### Signal length, *q*

If *q* = 0, then signals persist until the end of a trial; the optimal strategy in this case would be to only choose *strong* towards the end of a trial (taking account of how much total information it is ideal to collect). On the other hand, if *q* = 1, then a signal only ever lasts a single time step, and the cost of choosing *strong* even at a single time step outweighs the benefit in terms of enhanced detection (Fig. 8B, left). Between these two extremes, we observe that initially, as *q* increases, the occupancy of the *strong* attentional state increases, peaking at around *q* = 0.3; beyond this, the occupancy rate decreases, until at *q* ≈ 0.8, it is no longer worthwhile engaging the *strong* attentional state at all. This ‘breaking point’, where it is no longer worth paying the cost of *strong* attention, then marks where the performances of the optimal and always-*weak* policies converge (Fig. 8B, middle and right).

##### Information quality, *σ*_*s*_

To be worthwhile to choose *strong*, the informational benefit of this state must outweigh its cost. As we increase *σ*_*s*_ towards the value of *σ*_*w*_, it becomes less worthwhile to choose *strong*, until there is another breaking point (at *σ*_*s*_ ≈ 0.6) where it is no longer worthwhile at all (Fig. 8C, left). Again, this leads to a convergence in performance between the optimal and always-*weak* policies (Figs 8C, middle and right).

##### Attention costs, *κ* and *ϵ*

Very similar to the pattern of results seen when reducing the quality of information via *σ*_*s*_, as we increase the cost of a *weak* → *strong* shift (i.e., increase |*κ*|), there is again a point (|*κ*| ≈ 0.15) at which this cost is no longer worth paying (Fig. 8D, left), and we observe convergence of optimal and always-*weak* policies (Fig. 8D, middle and right).

For *κ* fixed, as we gradually make the cost of staying in the *strong* state more expensive (i.e., increase |*ϵ*|), we also observe a decrease in the willingness to occupy this state (Fig. 8E, left). At |*ϵ*| = |*κ*| = 0.1, where there is no benefit to maintaining the *strong* attentional state, it is still worthwhile to choose *strong*, but only for a single time step just before the decision is made (not shown).

### 3.2 ‘Optogenetic’ manipulations

Gritton et al. (2016) used optogenetics to either stimulate or suppress cholinergic activity while mice performed a sustained attention task (SAT); separate experiments focused either on cholinergic cell bodies in the basal forebrain (BF) or cholinergic terminals in the right medial prefrontal cortex (mPFC). They reported a rich pattern of results (Fig. 9A). On signals trials, while BF activation led to an increase in hits for shorter signals, and BF suppression led to a decrease in hits for longer signals, no significant effect on performance was found for mPFC manipulations. On non-signal trials, activation of either area led to an increase in false alarms, while suppression of BF had no effect on false alarms.

Figure 9B shows the effect on false alarms and hits when Gritton et al. activated (‘act.’) or suppressed (‘supp.’) the activity of ACh cells in BF. (Gritton et al. examined the effect of photoactivation/photoinhibition at different intensities; for simplicity, here we just re-plot the results for the most intense stimulation values: 21–25mW for photoactivation, and 11–14mW for photoinhibition.)

Despite the relative simplicity of our model, we were inspired by these results to measure the effect of certain manipulations. In the usual case we treated so far, it is assumed that the actual and believed qualities of the signal are the same. That is, for example, if the present choice is to attend *strongly* (*a*_*n*_ = *strong*), then the instantiation of *O*_*n*_ is indeed generated from 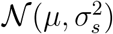, where *μ* depends on whether a signal is in fact present (*μ* = 1) or not (*μ* = 0), and inference proceeds on this basis (i.e., based on *𝒫*(*O*_*n*_|*X*_*n*_, *a*_*n*_ = *strong*)).

However, as well as considering different cases where this assumption is met, we can study cases where it breaks down, so that there is a mismatch between the actual and believed quality of the signal. We report results for the default set of model parameters (cf. Section 3.1.1), but the qualitative pattern of results holds for a broad range of model parametrizations. Figure 9C (left) shows the ‘matched’ case: observations are generated according to the optimal (‘control’), always-*strong* (‘act.’) or always-*weak* (‘supp.’) attentional policies, and the effect on inference is consistent/matched to the policy. As expected, hits increase (marginally) under always-*strong* and decrease under always-*weak*, while false alarms remain essentially the same (tiny decrease) and increase, respectively (cf. the hits and false alarms at *p*_1_ = 0.5 in Fig. 8A, middle). Modulo the relative sizes of these effects, this is as expected: more precise observations lead to better performance — and vice versa for less precise observations — and this is apparent in both hits and false alarms.

One type of mismatch we can consider is when observations are treated during inference as being generated under the optimal attentional policy, but are in fact generated under an always-*strong* (‘act.’) or always-*weak* (‘supp.’) policy (Fig. 9C, middle, ‘mismatch obs.’). This leads to a slight inflation of false alarms for the latter case, but otherwise yields the same qualitative pattern as the matched condition.

Another type of mismatch is when observations are indeed generated under the optimal attentional policy, but treated as being generated under always-*strong* (‘act.’) or always-*weak* (‘supp.’) policies (Fig. 9C, right, ‘mismatch inf.’). Interestingly, what we see here is qualitatively aligned with the BF findings of Gritton et al. (2016) (cf. Fig. 9B): hit rates are increased and decreased under ‘activation’ and ‘suppression’, but so too does ‘activation’ *increase* the false alarm rate, and ‘suppression’ (to a lesser degree) *decrease* the false alarm rate.

The fact that the latter, and not the former, type of mismatch captures the qualitative trend of results is notable as this allows us to decompose the putative three roles that ACh plays: in order for the (i) increase in the signal-to-noise ratio of the input to have an effect, the impact of the bottom-up information concerned has to be increased relative to prior expectations — something that can happen through (ii) boosting the strength of this signal, (iii) the suppression of priors, or both. The mismatch more consistent with the data (the rightmost plot of figure 9C) implicates (ii) and/or (iii), suggesting that this might be the dominant effect of the ACh manipulations. False alarms and hits both go up with BF activation, since the prior, which would be suppressed, implements the default assumption in this mode of signal processing that the signal is not present. Suppressing ACh boosts the relative effect of the prior, thus decreasing hits. The effect on false alarms is harder to assess, not the least because of a floor effect. That activating ACh just in the mPFC does not increase hits could be because the prior is already correctly incorporated in the sensory processing hierarchy, implying that hits cannot be substantially rescued at the level of the mPFC.

### 3.3 Multiple trials

So far, we have considered optimal control of attention when the optimization problem is to maximize performance for a single trial. In the single-trial case, it never makes sense to choose *strong* in the decision state, since one would be paying a cost for no advantage, informational or otherwise. In this section, we consider the optimal policy when future trials are also taken into account. We show that it can then make sense to maintain a *strong* attentional state into the next trial in order to avoid the future possible cost of a *weak* → *strong* shift, and suggest this may help explain a sequential effect in ACh signalling.

#### 3.3.1 Maintaining *strong* attention across trials

We keep the same, ‘default’ parameters as in the frequent signal, single-trial case, but we now solve for a policy that takes into account future trials; in particular, we solve for the policy that maximizes the average reward rate (see Methods).

The optimal policies in the *weak* and *strong* attentional states are essentially identical to the single-trial case; the only exception is that if one occupies the *strong* attentional state at the time of decision, then it is optimal to choose *strong* for all belief states (not shown) — and thereby start the next trial in the *strong* attentional state. This is despite the fact that there is no information to be gained in the decision state, and the following *N*_0_ steps of the next trial are known to be spent in the pre-signal state; the only reason to maintain *strong* into the next trial is because this is cheaper than paying the cost of a subsequent *weak* → *strong* shift.

Correspondingly, when examining average trajectories, one sees that on average, attention never completely reverts *strong* → *weak* at the end of a trial (Fig. 10). Furthermore, for a signal trial, the later the signal’s arrival, the more likely it is that *strong* attention is maintained into the next trial (Fig. 11).

#### 3.3.2 Sequential effects and ACh

Howe et al. (2013) measured ACh release in right mPFC while rats performed the SAT. They reported that ACh transients were only present on signal trials in which an animal correctly reported a signal (i.e., a hit trial) and not on correct rejection or miss trials; false alarm trials were not analyzed due to the comparatively low number of such trials (≈ 16% of non-signal trials). Furthermore, Howe et al. reported that a significant ACh increase occurred on a hit trial only when a non-signal was reported on the previous trial (‘incongruent hit’), while no such increase was observed on a hit trial if the previous trial was also a hit trial (‘consecutive hit’; Fig. 12A, left). A nonsignificant trend for a decline in signal during an ‘incongruent correct rejection’ (i.e., correct rejection when the previous trial was a hit trial) was also reported (Fig. 12B, left).

Dayan (2012b) suggested that if an initial detection of a signal establishes a detection-oriented task set that lasts across trials, then there would be little unexpected uncertainty when a (detected) signal follows (detected) signal — and so ACh release would not be expected during these consecutive hits (Fig. 12C; see also Hasselmo & Sarter, 2011). Implicit here is the additional requirement that *strong* → *weak* shifts do sometimes occur, thereby temporarily deactivating the detection-oriented task set — otherwise, there would be no interesting dynamics to observe at all.

In terms of our model, attentional dynamics — and, by hypothesis, so too ACh dynamics — that are similar to the reported pattern of ACh activation will arise in the model if the following are satisfied: (i) a *weak* → *strong* shift is more likely to occur on a signal trial than a non-signal trial (since an increase in ACh was only observed during signal trials); (ii) a *strong* attentional state is at least sometimes maintained into the next trial following a hit trial (since no increase in ACh was observed for consecutive hits); and (iii) *strong* → *weak* shifts do sometimes occur.

We already saw that a *weak* → *strong* shift is more likely to occur on a signal trial than a non-signal trial in the model if signals are expected to be rare (i.e., *p*_1_ ≪ 0.5; Fig. 7). We also saw a regime in which *strong* attention is maintained across trials (Fig. 10). What we have perhaps not examined so closely is the issue of when disengagement (i.e., a *strong* → *weak* shift) occurs. One might draw a distinction between passive and active mechanisms of disengagement: *active* disengagement would be mandated by the optimal policy (responding to the relative costs and benefits); while *passive* disengagement would not be so mandated, but rather occur sporadically, somewhat like a ‘lapse’ process, or perhaps even associated with the sort of spontaneous fluctuations in attention apparent in longer-running vigilance tasks (Fortenbaugh et al., 2017; Gilden & Wilson, 1995). The parameter *δ* in the model was introduced principally to allow for the latter possibility: even if the intention is to begin the next trial in the *strong* attentional state, one in fact begins in the *weak* state with probability *δ*.

Here, we focus on what is arguably the more interesting case — active disengagement. One possibility, amongst others (see Discussion), arises if we relax the very strong assumption that an agent knows the true parametrization of the task. For example, the fact that the true probability of a signal trial is *p*_1_ = 0.5, and is fixed at this level, is not a given. The signal probability needs to be estimated across trials, and may or may not be assumed to be stationary. Since the optimal policy depends on the estimated signal probability, if the latter changes, then so too should the policy (clearly, the same logic applies if there are nonstationarities in other model quantities, such as increases in cost with fatigue, etc.).

Consider the case where the estimated signal probability 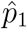 tends to be low, but changes over time with trial-to-trial feedback. Furthermore, assume that for relatively high estimates, it is optimal to maintain *strong* attentional state across trials, while for lower estimates it is not. Since a signal trial will lead to an increase in 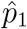, the occurrence of a signal trial will encourage maintenance of the *strong* attentional state if it is occupied; conversely, a non-signal trial will lead to a decrease 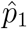, thus discouraging maintenance of a *strong* attentional state.

In Figure 13, we show an example of the evolution of beliefs, attention, and estimated signal probability over 6 trials. For simplicity, we assume that 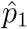 switches between only two (low) values: a higher probability 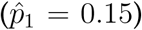 and a lower probability 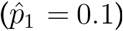. Changes between these estimates are prompted in two ways. The first way is that if 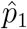 is currently low, and a *weak* → *strong* shift occurs during the trial, then this shift immediately prompts 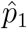 to be increased during the trial, and this lasts at least until feedback at the end of the trial. This can be justified by considering that a *weak* → *strong* shift always indicates a suspicion that a signal may be present, whereas a *strong* → *weak* shift does not necessarily indicate a belief that a signal is absent (indeed, we saw above that switches of the latter type are more associated with certainty that a signal has occurred). The second way is in response to feedback (i.e., reward vs. no reward) at the end of each trial, which, at least from the experimenter’s point of view, unambiguously indicates whether a signal was or was not in fact present in the trial. Therefore, according to this latter assumption, we deterministically set 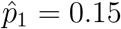 after a signal trial, and 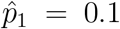 after a non-signal trial. A change in 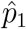 yields an immediate change in policy, and, in our chosen example, *strong* is always maintained across trials when 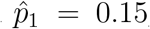, but not when 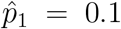 (parameters otherwise set as follows: *q* = 0.2, *σ*_*w*_ = 1, *σ*_*s*_ = 0.5, *κ* = −0.1, *ϵ* = −0.005, *δ* = 0.001).

**Figure 13:**
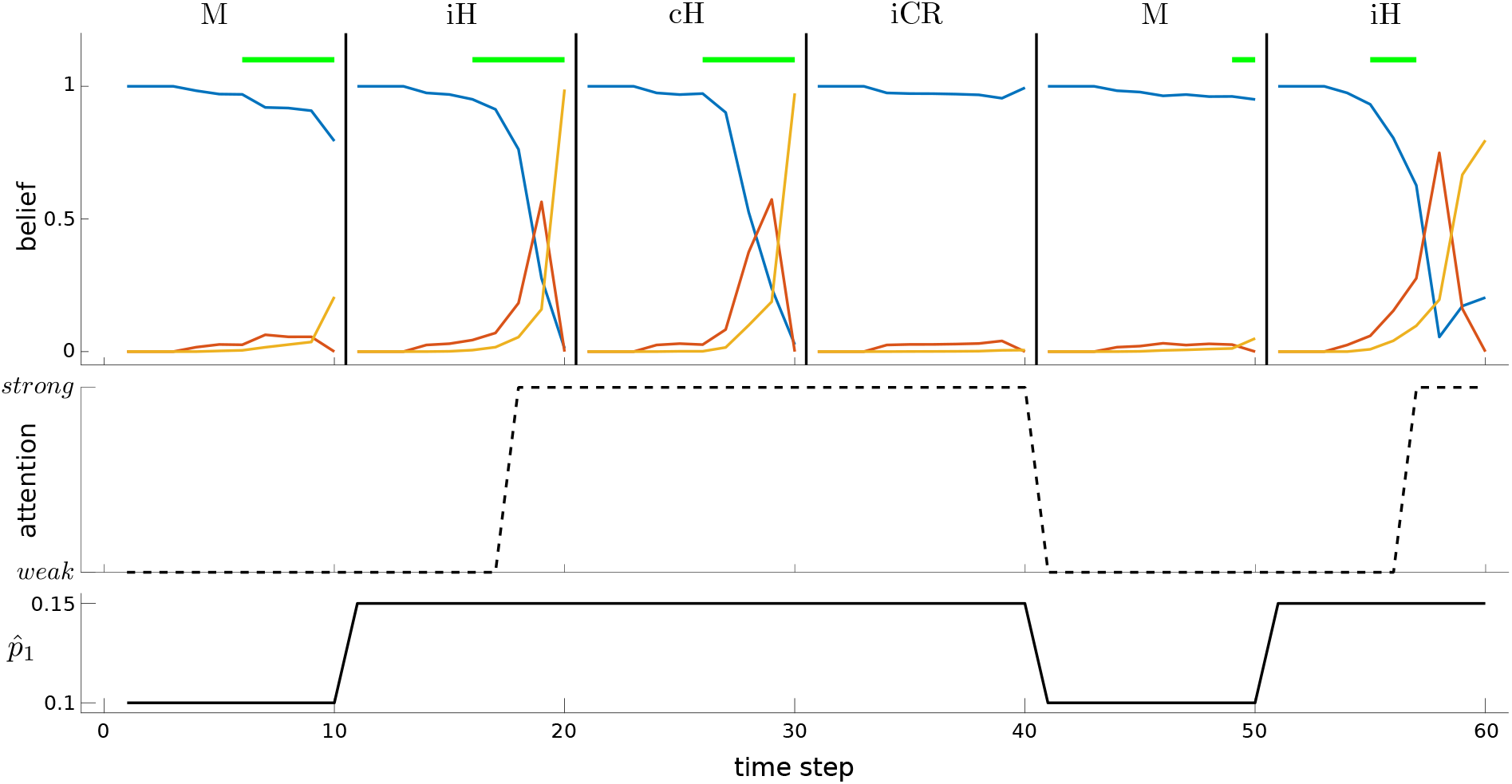
Attentional shifts with nonstationary estimates of signal probability. See main text for detailed description. Top: beliefs over time for each trial (vertical black lines indicate trial boundaries); horizontal green lines indicate a signal; a trial can be classified as a miss (M), hit (H), or correct rejection (CR; there are no false alarms in this example), and as either ‘congruent’ (c) or ‘incongruent’ (i) depending on what occurred on the previous trial. Middle: changes in attentional state. Bottom: changes in estimated probability of a signal, 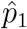.

The first trial is a miss (M) trial (i.e., there was a signal, but a non-signal was reported), but the feedback indicates that a signal was in fact present, and so 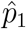 is increased. Trial 2 is also a signal trial, but during this trial there was a *weak* → *strong* shift, and the response was correct (hit); this is an incongruent hit (iH). Since 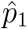 is already high, the *strong* attentional state is maintained into Trial 3. This trial is also a signal trial, and the response is correct (i.e., congruent hit (cH)) — but crucially, since already in the *strong* state, there is no change in attentional state. Trial 4 is a (incongruent) correct rejection (iCR) trial, and the lack of a signal leads to a reversion of 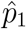 to a lower value. Trial 5 is a miss trial, leading 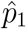 to be increased at its end. Finally, Trial 6 sees a *weak* → *strong* shift, and is a hit.

The point of this example is to show one way in which incongruent hits can be associated with *weak* → *strong* shifts (i.e., relative ACh activation), while congruent hits can be associated with no change in attentional state (i.e., no change in ACh). Indeed, if we run a large number of trials, and treat attentional/ACh dynamics in the model similarly to how Howe et al. treated their ACh data (i.e., baseline-corrected and aligned), we find a similar difference between congruent vs. incongruent hits (Fig. 12A, right).

Interestingly, in the model, we see a rather large negative change at the end of an incongruent correct rejection trial (Fig. 12B, right). This reflects a *strong* → *weak* shift: the previous trial is a hit trial, on which the *strong* attentional state is likely to be occupied and maintained into the next trial (since 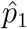 will be high); the current trial is a non-signal trial, during which *strong* tends to be maintained throughout, but at its end, feedback leads to a decrease in 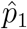 (since there was indeed no signal), leading to a *strong* → *weak* shift (i.e., relative decrease in attention/ACh).

Of course, Howe et al. (2013) observed only a non-significant, and apparently transitory, trend towards a decrease in ACh on incongruent correct rejections, and so this latter pattern of model results is, strictly speaking, a misprediction. Nevertheless, we think it perhaps interesting to consider, since it is the mirror of the case for hit trials: i.e., incongruent hits are associated with a *weak* → *strong* shift because attention is likely to start *weak* (because of preceding trial type) and be activated during the course of the current, signal trial; incongruent correct rejections are associated with a *strong* → *weak* shift because attention is likely to start *strong* (due to preceding trial type) and be deactivated at the end of the current, non-signal trial (Fig. 12D). Our misprediction is likely to be influenced by a particular simplifying assumption that we made concerning when 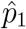 is updated, namely only at the end of the trial. Thus, in the simulation results in Figure 12A, the incongruent hit signal represents averaging over *weak* → *strong* shifts occurring on different time steps (due to different possible onsets of the signal). By comparison, in Figure 12B, the incongruent correct rejection signal averages over *strong* → *weak* shifts that all occur at the same time (i.e., trial end). The latter thus leads to a much stronger (and consequently mispredicted) decrease in attention/ACh. In this light, it would certainly be interesting to consider more realistic assumptions about ongoing estimation of 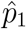, as well as other modifications of the model.

## 4 Discussion

### Summary

We considered vigilance, or sustained attention, from the point of view of an optimal control problem in which a series of choices of internal, attentional action are made, and where these internal actions may accrue both costs and benefits. In particular, inspired by the sustained attention task (SAT; McGaughy & Sarter, 1995) developed for rodents, we considered the control of a binary attentional state in an abstract model of the SAT involving trials in which *strong* attention yields better sensory information than *weak*, but incurs costs. We analysed how optimal allocation of attention (i.e., the choice of which attentional state to occupy when) depends on signal characteristics (whether a trial is likely to contain a signal, and if so, when it is likely to be on), the relative qualities of information in the different attentional states, and the costs associated with paying *strong* attention. We also considered what happens when the actual and expected signal properties do or do not match, and possible relationships with the effects on detection performance observed in an optogenetic study.

In the multi-trial case, we saw that optimal allocation of attention also takes into account performance in future trials; it can be optimal to pay the costs of maintaining *strong* attention into the next trial, not because of any immediate informational benefit — there is none — but in order to avoid the future cost of a *weak* → *strong* shift. We demonstrated that this can lead to trial history effects when *weak* → *strong* shifts occur, and suggested that such sequential effects may help explain when cortical ACh transients have been observed to occur in the SAT.

### Elective attention

We treated the elective allocation of attention as a characteristic form of internal action (Dayan, 2012a). Such internal actions can be considered on an equivalent footing with external actions, potentially coming with actual costs associated with the active deployment of neural resources, opportunity costs associated with the unavailability of those resources for other tasks (which we did not consider; see Boureau et al., 2015), and, in our case, well-defined benefits in terms of increasing the signal-to-noise ratio (SNR) of external inputs. Our calculations are examples of the expected value of control (EVC; Shenhav et al., 2013, 2017), and may involve similar psychological and neural considerations. Indeed, it has been observed that rewarding human subjects for their performance on sustained attention tasks can improve their momentary performance (Engelmann, Damaraju, Padmala, & Pessoa, 2009; Esterman, Reagan, Liu, Turner, & DeGutis, 2014), consistent with the proactive engagement of various cortical attention networks (Esterman, Poole, Liu, & DeGutis, 2017). In such tasks, there is often an overall decrement in vigilance over the course of a session, which is also subject to varied motivational effects (e.g., Esterman et al., 2016, 2014; Massar, Lim, Sasmita, & Chee, 2016). We did not address vigilance decrement in the current study, particularly since this is not a reliable feature of SAT performance (McGaughy et al., 1996; McGaughy & Sarter, 1995; M. Sarter, personal communication, July 2022). However, given the prominent place of vigilance decrement in the sustained attention literature, and recent intriguing reports of dissociable influences of motivational manipulations on overall performance versus performance decrements over time (Esterman et al., 2014), this would certainly be a salient target for future work.

From the perspective of a general POMDP (Kaelbling et al., 1998), the way that information-gathering is instrumental for reward-gathering (or cost-avoiding) is completely standard. It also arises in the context of sequential probability ratio tests or drift diffusion decision-making (Gold & Shadlen, 2007; Ratcliff, 1978), where the benefits of gaining new information about an uncertain external stimulus are balanced against the opportunity cost of time. In economics, considerations such as ours are associated with the framework of rational inattention (Caplin & Dean, 2015; Hébert & Woodford, 2017, 2019; Sims, 2003) in which agents are charged for the mutual information between internal representations and external signals, and so, like our agent, opt to remain partially ignorant. Recently, Mikhael, Lai, and Gershman (2021) adapted the rational inattention framework to the case that the neuromodulator dopamine (DA), by reporting the average reward rate in an environment (Niv, 2007), controls the fidelity of the internal representation of the external information (for similar lines of thought, see, e.g., Cools, 2019; Manohar et al., 2015; Westbrook & Braver, 2016; Westbrook et al., 2020). In our terms, this suggests a rich relationship between DA, as the medium of expected value, and ACh, as the medium of arousal and attentional improvement. For example, St. Peters, Demeter, Lustig, Bruno, and Sarter (2011) provided evidence that NMDA-mediated stimulation of the shell (though not the core) of the nucleus accumbens — a principal target of dopaminergic neurons in the ventral tegmental area — can, via activation of cholinergic cortical projections, counteract distractor-induced impairments of performance in the SAT (though interestingly, performance was not improved by such stimulation in the absence of distractors).

We assumed just two attentional states, *strong* and *weak*, which has the practical benefit of facilitating analysis, but is also consistent with interpretations of behaviour in the SAT in terms of shifts between a small number of task-sets or ‘modes’, such as ‘default’ and ‘detection’ modes (Hasselmo & Sarter, 2011; see also below). Nevertheless, the model could readily be extended to allow for different degrees of attention incurring different costs (as, for instance, in the graded information-based costs in rational inattention; Hébert & Woodford, 2017, 2019), thereby permitting more fine-grained control of the SNR.

### Heterogeneous attentional deployment

Particularly when engaging and/or maintaining attention is expensive, it would often be better to have either *strong* or *weak* attention for the whole trial — and, indeed, this is conventional for signal detection theory (Green & Swets, 1966). Here, we investigated the more subtle case when the agent collects sufficient information to conclude that it becomes worth engaging attention part-way through a trial, leading to a variable rate of information gathering. The work therefore has natural links with previous studies that consider optimal integration for (uncontrolled) time-varying reliability of evidence (e.g., Cisek et al., 2009; Drugowitsch et al., 2014) and control of visual fixations (e.g., Jang et al., 2021; Tavares, Perona, & Rangel, 2017). The structure of the task considered here, in which the signal never comes on for a set period after initiation and is more likely to be on towards the end of a trial, implies that the participants should engage in a form of temporal orienting of attention (Nobre & Rohenkohl, 2014) — most obviously, refusing to pay attention immediately after a trial starts. Of course, such temporal considerations are subject to the vagaries of interval timing (Gibbon, 1977), which would rapidly make this imprecise over the timescale of several seconds. In our case, this conditional engagement of attention arises under a somewhat particular set of parameter values in which, very crudely, the default operating regime is to assume the absence of a signal (due to its actual/assumed low probability of occurrence in a trial and/or temporal sparsity within a trial), but if a signal is suspected, then it is worth confirming with the higher precision information available following *strong* attentional engagement. False alarms could then arise from a form of lapse process or ‘trembling hand’, which, for convenience, we did not model. Altogether, this makes the task involve a form of vigilance (Broadbent, 1971; Luce & Green, 1974; McGaughy et al., 1996). Note that in the regime we assumed in relation to Howe et al. (2013), although a signal is as likely to appear as not on any given trial, we set the *assumed* probability of a signal in the model to be much lower, which is why *strong* attention was more likely to be engaged on signal trials. There was therefore a mismatch between the ‘true’ generative model (of the experimenter) and the model ascribed to the agent. While this was somewhat pragmatic in our hands, we suspect such mismatches to be not uncommon. In the particular case of the SAT, subjects ought, if they understand that the feedback at the end of each trial reliably indicates whether a signal was present, to apprehend that the long-run relative frequency of a signal trial is approximately one half — but only then if they also (correctly) believe that this relative frequency is fixed.

Other cases of heterogeneous attentional engagement have been studied when external information signals that the environment has switched into a markedly different state (e.g., going from safe for foraging to dangerous; Lloyd & Dayan, 2018). This has been considered as a sort of interrupt signal that involves norepinephrine (NE) rather than ACh (Bouret & Sara, 2005; Dayan & Yu, 2006). The case of danger also involves arousal, but more a form of cognitive reset and task-switching from unexpected uncertainty than a controlled engagement of focus within a single task arising from expected uncertainty (Yu & Dayan, 2005). The task that Dayan and Yu (2006) used to examine phasic NE (Clayton, Rajkowski, Cohen, & Aston-Jones, 2004) also involved signal detection — but NE was treated purely as a read-out mechanism for a state change in the external world, rather than having an effect on stimulus processing as here. It would certainly be intriguing to also measure the involvement of NE in the task we considered.

Shea-Brown, Gilzenrat, and Cohen (2008) previously considered the involvement of NE in a signal detection task. For them, NE release also depended on excursions beyond a threshold of certainty about the presence of a signal. However, NE played the role in their model of destabilizing the dynamics of a downstream decision-making network so that it stopped integrating new information, and instead evolved quickly to a boundary in state space at which an action would be initiated. ACh plays a radically different role in our model by regulating — rather than curtailing — ongoing integration. One important consideration raised by Shea-Brown et al. (2008) is the speed of action of the neuromodulator. NE takes its effect in cortex rather slowly — at least around 100ms after electrical stimulation of the source nucleus, the locus coeruleus (Waterhouse, Moises, & Woodward, 1998). A similar speed of action for ACh (Pinto et al., 2013) might suggest that its phasic engagement might be too sluggish to improve the processing of short signals (the shortest were 25ms and 50ms in Howe et al., 2013), though conceivably fast enough to affect neural activity prompted by the signal that outlasts the signal itself, even at relatively early stages of vision (e.g., Offen, Schluppeck, & Heeger, 2009).

On a rather different timescale are the fluctuations in attention typically observed over the course of minutes to hours in the long-run vigilance tasks often used to interrogate sustained attention in humans (reviewed, for instance, in Esterman & Rothlein, 2019; Fortenbaugh et al., 2017; Gilden & Wilson, 1995). These are also associated with NE (e.g., slow fluctuations in tonic activity, which are anticorrelated with phasic responses to task-relevant stimuli; Aston-Jones & Cohen, 2005; Howells, Stein, & Russell, 2012). Providing an integrated understanding of the effects across these various timescales, perhaps including their contribution, if any, to spontaneous *strong* → *weak* attentional transitions, is an important task for the future.

### Acetylcholine

We designed and illustrated our account in the light of the sustained attention task (SAT) of Sarter and colleagues (Gritton et al., 2016; Howe et al., 2013; McGaughy & Sarter, 1995). However, given the complexity of the paradigm, we sought a qualitative rather than quantitative match with their results. ACh is known to play a causal role in this task because of the effects of pharmacological and optogenetic suppression and stimulation (Gritton et al., 2016; McGaughy et al., 1996). In our information-processing operating regime, the default is that the signal is absent and ACh is involved in the choice to gather sufficient information to refute this presumption. Thus, it is no surprise that the main effect of suppressing ACh is to reduce detection (i.e., reduce hits). That stimulating the BF increases detection can be explained in the same way — allowing higher-quality information about signal presence to be collected — particularly aiding detection of signals that are not normally able to benefit from ACh release in virtue of being too short. It is notable that activation of ACh axon terminals in right mPFC had no effect on signal trials (Gritton et al., 2016).

The observation that stimulating the BF also increases false alarms (Gritton et al., 2016) is harder to reconcile with the simple view that the release simply increases the quality of sensory information. One possibility, hinted at by the account we gave in which there was a discrepancy between the assumed and actual information, is to note that there are two ways that information integration can be affected in the light of a changed relative SNR in the external input versus the internal default. One is for there to be extra activity in the representation of that input, such as extra spikes associated with the signal that can overwhelm meagre activity in the integrator representing signal absence (Ma, Beck, Latham, & Pouget, 2006). The other is for recurrent interactions in the integrator to be suppressed in favour of the external signal (Gil, Connors, & Amitai, 1997; Hasselmo & Sarter, 2011; Hsieh, Cruikshank, & Metherate, 2000; Kimura, Fukuda, & Tsumoto, 1999). To the extent that the latter happens when the external signal is actually not of any better quality, the more likely it is that a false alarm could arise on a non-signal trial. It is likely that both mechanisms are engaged, and over multiple layers of cortical inference about the signal; disturbance of the delicate balance between internal expectations of signal unlikeliness and external signal and noise would lead to false alarms. Unfortunately, Gritton et al. (2016) did not report whether trial sequence also affected the consequences of optogenetic manipulations, as one might suspect given the previous report of Howe et al. (2013).

In contrast to the effects of stimulating the cell bodies of cholinergic neurons in the BF, Gritton et al. (2016) found that activation of their axons in right mPFC in non-signal trials led to an increase in false alarms, but activation in signal trials did not improve detection. The authors speculated that activation of ACh axon terminals in mPFC may be sufficient to modulate behaviour under circumstances where endogenous ACh transients are not expected (i.e., non-signal trials), but insufficient to modulate behaviour in circumstances where endogenous ACh transients are expected (i.e., signal trials) — BF activation/suppression, by modulating ACh release more widely, may exert a stronger influence. Unfortunately, technical limitations meant that they did not record cholinergic transients concurrently with optogenetic manipulations. In our terms, stimulation in PFC might only affect the suppression of the default integration — and so mainly lead to problems with false alarms.

Note that the authors of the original study have a different interpretation of the ACh release, at least in PFC. That is, Howe et al. (2013) interpret their findings in terms of a mechanism supporting shifts from ‘externally-directed’ attention, which in this case involves sensory monitoring for the possible appearance of a signal, to ‘internally-directed’ attention, here involving the retrieval and (re)activation of signal-associated response rules when a signal is detected (cf. Chun, Golomb, & Turk-Browne, 2011). Thus, they consider ACh to be engaged following full-blown detection, but to be involved in the correct response rather than in refined signal processing. Indeed, Sarter, Hasselmo, Bruno, and Givens (2005) cite results from Holley and Sarter (1995) showing that selective cholinergic deafferentiation of primary and secondary visual cortices does not alter animals’ SAT performance. Of course, one of the beneficial characteristics of the task is that signal and no signal have the same response demands (just requiring different levers to be pressed) — adding some complexity to this account.

Pinto et al. (2013) used optogenetics to activate or inactivate ACh neurons in BF in awake mice selectively while they performed a visual go/no-go task involving discriminating between a vertical (target) and horizontal (non-target) drifting grating; task difficulty was manipulated by adjusting stimulus contrast. They found that ACh activation led to enhanced discriminability at all contrast levels, and this was due to a selective increase in the hit rate (there was no significant change in false alarm rate). Activation of cholinergic axons in V1 also led to enhanced discriminability attributable to increased hits, though not at the highest contrast. This could be consistent with enhanced processing of the input stimuli if the targets were more temporally dense (indeed, in this paradigm, 1s after an initial cue, a drifting grating was presented for 4s; the intertrial interval was 3s), so that there was no equivalent default of ‘nogo’ as in Gritton et al. (2016); Howe et al. (2013). Inactivation of ACh cells in BF reduced sensitivity at all contrast levels. However, this may be due to an increase in false alarm rate, rather than a decrease in hits (L. Pinto, personal communication, December 2021), which could be harder to explain.

Pinto et al. primarily targeted the nucleus basalis, though they note the possibility that ACh neurons in other BF nuclei may also have been activated by optogenetic stimulation; retrograde tracing revealed that cholinergic neurons throughout the basal forebrain project their axons to V1. Dual retrograde tracing revealed very few ACh neurons projected to both V1 and mPFC (5%), or to both V1 and primary auditory cortex (6%), while a slightly higher number projected to both V1 and neighbouring higher visual areas (13%).

However, we should note that some studies of the activity of ACh neurons do not find the sorts of responses that our analysis would expect. For instance, Hangya, Ranade, Lorenc, and Kepecs (2015) (see also Laszlovszky et al., 2020) recorded from optogenetically-identified ACh neurons in the BF of mice during an auditory task, in which animals had to respond to one tone with a lick to gain a water reward (‘go’ stimulus), and withhold licking in response to a differently pitched tone to avoid an airpuff punishment (‘no-go’ stimulus); to manipulate difficulty, the loudness of tones was also varied. Hangya et al. reported that most recorded ACh neurons were activated by both reward and punishment delivery. Neural responses to reward were modulated by the intensity of the preceding tone, so that the strongest neural response was observed following the quietest tone, when the animal may be least certain of the outcome — consistent with the coding of a degree of ‘reinforcement surprise’; neural responses to punishment showed no such modulation by stimulus strength. Only 2/34 ACh neurons showed attentional modulation, operationalized as activity before stimulus onset that predicts RT or accuracy. Although reinforcement surprise is a key driver of expected uncertainty, consistent with the view of ACh reported above (Yu & Dayan, 2005), the lack of attentional effect in this study is more puzzling. The more recent study of Laszlovszky et al. (2020), which reports a greater degree of heterogeneity in ACh responses, and evidence about local control of release of ACh (Parikh, Man, Decker, & Sarter, 2008) may help resolve such puzzles. It may also be important, as with Pinto et al. (2013), to consider distinctions between the paradigm used by Hangya et al. (2015)/Laszlovszky et al. (2020) compared to the SAT, such as differences in stimuli, timing, and response requirements (e.g., go/no-go lick vs. press left/right lever). Further clues may come from studies of the action of ACh at a detailed, implementational, level such as, for instance, recent modelling work that uses excitatory-inhibitory networks to explore how ACh may promote local gamma oscillations and theta-gamma coupling (Lu, Sarter, Zochowski, & Booth, 2020; Yang et al., 2021).

Along with these effects within a single trial is the evidence about the engagement of ACh when considering the sequential structure of trials (Howe et al., 2013). Here, perhaps the most surprising finding is that ACh is particularly activated on a signal trial when the previous trial either did not involve a signal, or it did but the signal was missed. We interpret this vivification of ACh as being associated with the deployment of attention. Indeed, the observed ACh transients are thought to be generated *within* PFC (i.e., local release), based on interactions between thalamic glutamatergic afferents and heteroreceptor-mediated regulation of cholinergic terminals, rather than reflecting phasic activity of BF ACh neurons (Parikh et al., 2008). Howe et al. speculate that during consecutive hits, ACh activity associated with the first (incongruent) hit may induce persistent spiking and continuing release of ACh that helps maintain the activated (signal-oriented) task set. Decay of the latter activation would then lead to a return to the predominant monitoring state. One prediction we can make related to sequentiality is that the short signals will be better detected when attention was already engaged at the beginning of the trial on which they occur, based on the events on the previous trial.

Our account of why these sequential effects arises is rather fragile, and depends on two key assumptions. The first is that once attention is engaged in a trial, it is worth keeping it engaged even after the agent is statistically convinced (or has reported) that the signal has been present. This depends on the cost of maintaining *strong* attention being much less than a *weak* → *strong* switch, and also being outweighed by the benefit of starting a new trial with *strong* attention engaged (including, for instance, the excess probability of correctly detecting a very short signal, or an immediate change in the assumed likelihood that trials will include a signal). The second assumption is that attention will sometimes revert to *weak*, thus requiring another *weak* → *strong* shift; in our parameter regime, these latter shifts will happen preferentially on signal trials. We focused on an active mechanism of disengagement (i.e., *strong* → *weak* shift) arising from fluctuating estimates in signal probability but, as noted, changes in other parameters could also play a role; utilities, costs, signal-to-noise ratios could all plausibly fluctuate over the course of a trial as an animal’s state of satiety and/or fatigue varies (e.g., Gergelyfi, Jacob, Olivier, & Zénon, 2015). The model could also accommodate passive shifts or sporadic distraction, or indeed the sort of ‘in-zone’ to ‘out of zone’ spontaneous fluctuations that happen in long-run vigilance tasks (Fortenbaugh et al., 2017; Gilden & Wilson, 1995), are affected by motivational factors (Esterman et al., 2016), and might be associated with NE (Aston-Jones & Cohen, 2005; Howells et al., 2012). The converse spontaneous increases in attention could be similarly modelled.

Our account followed Hasselmo and Sarter (2011) in treating *weak* and *strong* states as forms of task-sets or ‘modes’: a default response mode, in which the animal simply executes the most extensively-practiced response (i.e., presses the lever to report a non-signal, which is indeed the majority response) and a detection mode. The costs in switching between the two are then analogous to the switch costs that are prevalent in the task-switching literature (e.g., Monsell, 2003). We assumed that switch cost only accrues for the shift away from the default.

One important predecessor of the current work is by Atkinson (1963), who assumed that maintaining a high level of sensitivity would be costly, and so subjects would aim to adjust their sensitivity level dynamically to trade off any consequent decrement in performance against the minimization of this cost. He specified particular rules for updating, trial to trial, both the level of sensitivity and the decision rule (i.e., criterion), and noted that sequential effects in detection performance could arise from adjustments to either or both. Partly inspired by this earlier work, Gilden and Wilson (1995) studied ‘streakiness’ of performance (i.e., non-stationarity in correct vs. incorrect responses across trials) in signal detection tasks — akin to our discussion of maintained and lapsed attention. Gilden and Wilson interpreted their results as suggesting that the greater the attentional resources demanded in a task, the lower the level of streakiness; they also considered the possibility that the delay between trials is an important determiner of sequential dependence, where longer delays encourage greater independence between trials, though their evidence on this was mixed. We note that Sarter et al. (2016), in interpreting ACh transients observed during a task slightly simpler than the SAT (rats were trained to approach a port to retrieve a reward following a visual signal — see Parikh, Kozak, Martinez, & Sarter, 2007), suggest that the much longer ITI involved (90*±*30s) meant that ACh transients were reliably elicited when animals detected the signal — since animals would most likely disengage during the long ITI, meaning that any subsequent successful detection would be an ‘incongruent hit’ (and thus elicit ACh release). Gilden and Wilson (1995) also considered a number of models that would give rise to streaky outcomes, including a model of ‘intermittent attention’ with probabilistic transitions between a low effort state (with lower hit rate) and a high effort state (the conditional transition probabilities were assumed to be fixed). Similar such switching in engagement has been more recently studied in the context of animals solving a series of signal detection tasks (Ashwood et al., 2022; Weilnhammer, Stuke, Eckert, Standvoss, & Sterzer, 2021), and may relate to the fluctuations in attentional engagement found over the long run in human sustained attention tasks that we mentioned above (Esterman & Rothlein, 2019; Fortenbaugh et al., 2017; Howells et al., 2012).

Our task and analysis concerns a relatively simple signal — the turning on and off of a light of fixed intensity and location, and of variable duration (i.e., unimodal, and varying along one dimension) — and response (i.e., press one of two levers), and so is likely not ideal for addressing heterogeneity in cholinergic neuromodulation, let alone the substantial additional complexities of cortical control over attentional enhancement (e.g., Miller & Buschman, 2013) and its motivational sensitivities (Engelmann et al., 2009; Engelmann & Pessoa, 2007; Esterman et al., 2016, 2017). Heterogeneity in all the neuromodulatory systems is being intensively investigated (Lammel, Lim, & Malenka, 2014; Ren et al., 2018; Totah, Neves, Panzeri, Logothetis, & Eschenko, 2018), and ACh has long been identified as being more specific (Záborszky et al., 1985). This is appropriate to the extent that ACh reports on expected uncertainty, given that such expectations can be diverse.

## Conclusion

Stepping back from the details of this task and model, our simulations exemplify ways in which a neuromodulator might provide central regulation and co-ordination of the sort of radically distributed processing that otherwise occurs in the context of point-to-point, wired, connections.

## Acknowledgements

We are very grateful to Howard Gritton, Matt Howe, Lucas Pinto, and Martin Sarter for their helpful comments on previous versions of the manuscript. This work was supported by the Max Planck Society and the Alexander von Humboldt Foundation.

